# Energy-based Analysis of Biochemical Oscillators Using Bond Graphs and Linear Control Theory

**DOI:** 10.1101/2024.06.06.597695

**Authors:** Peter J. Gawthrop, Michael Pan

## Abstract

The bond graph approach has been recognised as a useful conceptual basis for understanding the behaviour of living entities modelled as a system with hierarchical interacting parts exchanging energy. One such behaviour is oscillation, which underpins many essential biological functions. In this paper, energy-based modelling of biochemical systems using the bond graph approach is combined with classical feedback control theory to give a novel approach to the analysis, and potentially synthesis, of biochemical oscillators. It is shown that oscillation is dependent on the interplay between *active* and *passive* feedback and this interplay is formalised using classical frequency-response analysis of feedback systems. In particular, the *phase margin* is suggested as a simple scalar indicator of the presence or absence of oscillations; it is shown how this indicator can be used to investigate the effect of both the structure and parameters of biochemical system on oscillation. It follows that the combination of classical feedback control theory and the bond graph approach to systems biology gives a novel analysis and design methodology for biochemical oscillators.

The approach is illustrated using an introductory example similar to the Goodwin oscillator, the Sel’kov model of Glycolytic Oscillations and the Repressilator.

## 1 Introduction

Oscillatory behaviour underpins many essential biological functions from circadian rhythms to cardiac pacemaking [1]. Oscillations of molecular species concentrations within the reaction networks of cells have been observed and analysed extensively using mathematical approaches over a long period [2–9]. For example, the Goodwin Oscillator has been recently discussed and placed in a broader research context by Gonze and Ruoff [10], the Glycolytic Oscillator of Sel’kov [11] is analysed by Keener and Sneyd [12] and a synthetic oscillator using DNA transcription, known as the repressilator, was introduced by Elowitz and Leibler [13] and discussed by Tyson et al. [14].

A key feature of biochemical oscillators is that the oscillation can only continue with a sustained supply of energy [15, 16]. This key feature can be included in the oscillator model using the *bond graph* approach [17–21] introduced into Systems Biology by Oster et al. [22, 23] and further developed by Gawthrop and Crampin [24]; an introduction to the approach is given by Gawthrop and Pan [25]. More generally, the bond graph approach has been recognised as a useful conceptual basis for understanding living entities as a system with hierarchical interacting parts exchanging energy [26, 27]. This paper sets biochemical oscillators within this conceptual framework using a modular hierarchical bond graph approach [28–31]. Oscillators have been considered in the bond graph context by Stramigioli and van Dijk [32], and a bond graph model of a repressilator [14] is developed by Pan et al. [16].

In the biochemical context, bond graphs imply the corresponding Chemical Reaction Networks (CRN) [33, 34] which are described by nonlinear ordinary differential equations (ODE). Nonlinear ODEs have an rich set of behaviours [35, 36]. One such behaviour is the *limit cycle* where the *n*_*x*_-dimensional system state *x* periodically moves along a closed trajectory in the *n*_*x*_ dimensional state space. Oscillations in biological systems are described by limit cycles [37–41] which therefore form the focus of this paper. Predicting the limit cycle behaviour of a nonlinear ODE is, in general, a difficult problem [35, 36]. However, in the special case of a second-order system where *n*_*x*_ = 2 (and so the state-space is planar), the presence or absence of limit cycles is predicted by the Poincaré-Bendixson theorem [35, 36]. In some cases, the behaviour of a high-order system can be approximated by a second-order system; in particular, the Liénard oscillator, a class of second-order systems representing non-linear oscillators, has been suggested as a universal model of biological oscillators [42].

Oscillations in biological systems are typically associated with *feedback* [38, 41]. The discipline of control engineering is concerned with feedback in general and feedback-induced limit cycles in particular [43–46]. A standard analysis tool is the *Describing Function* which extends the frequency domain approach to a limited class of systems comprising a linear dynamical system connected to a nonlinear non-dynamical system in a feedback configuration. Unfortunately, biological oscillators are not within this limited class and so the describing function cannot be used directly. Nevertheless, it is shown in this paper that the underlying philosophy, based on linear systems analysis using frequency response and the Bode diagram, can be used to elucidate the relationship between the physical system and its oscillatory properties.

“Approximation to a non-linear system by linearising at an equilibrium is an important and generally useful technique. If the geometrical nature of the equilibrium point can be settled in this way, the broad character of the phase diagram often becomes clear.” [35]. Moreover, analysis of a linear system is far easier than that of the underlying non-linear system and so, as will be illustrated here, it is possible to use the linearised model to rapidly sift though sets of system parameters guided by intuition based on the linearised system. Although a linearisation approach to nonlinear systems analysis cannot capture the full non-linear behaviour, it does allow the rich theory of linear feedback control design [47] to provide the initial stages of biochemical oscillator analysis and, potentially, *synthesis*. The resultant design may then be analysed and refined using nonlinear techniques such as the Hopf bifurcation method [12, 35, 36] and Lifting [48].

A number of previous approaches [5–9] consider the linearisation of biological oscillators to examine local stability. This paper extends the linearisation approach within the context of an energy-based model and includes frequency response analysis. The concepts of active and passive feedback have been used to analyse energy-based bond graph modelling of the feedback control of biomolecular systems [49]; these concepts have been repurposed in this paper. In particular, frequency domain analysis via the Bode diagram give insight into how the interplay between active and passive feedback gives rise to oscillations. The *root-locus diagram* is another analysis tool for linear feedback systems. It has been used in the context of biochemical oscillators previously [8]; here, the approach is extended and its advantages and limitations discussed.

Although standard control theory clearly distinguishes the concepts of controller and system (to be controlled), the distinction is not as clear in the context of biological systems. However, the concept of *physical-model-based control* [50–56] treats the controller as another physical system and thus is directly applicable in the context of biological systems. Coupled physical systems in general, and biological systems in particular, exhibit *retroactivity* [57–59], whereby components of biological systems change behaviour when coupled. This distinction between computational modularity and behavioural modularity has been explained by the bond graph approach [28]. As discussed previously [49], physical-model-based control explicitly takes account of the interactions between physical systems and thus implicitly acknowledges retroactivity. Moreover, such interactions are regarded as beneficial, rather than an unwanted artefact.

As shown previously [49], bond graph feedback can be partitioned into *active* feedback and *passive* feedback – the latter corresponding to retroactivity. Passive feedback has a generally stabilising effect which is often beneficial when stable system behaviour is required. However, in this paper it is shown that analysing the interplay between the active and passive feedback is crucial to understanding, and therefore creating, the conditions leading to oscillation.

Because the paper combines approaches from three fields – systems biology, bond graphs and control theory – there are three notational systems. To summarise, a generic species A has a bond graph representation **Ce**:**A** and the corresponding “signal” is the concentration *x*_*A*_. In a particular case, the generic species A can be instantiated as a particular species. The species *chemical potential* is denoted as *ϕ*_*A*_ and the species flux is denoted as *f*_*A*_.

The computation was performed using Python within Jupyter Lab [60], together with the bond graph package `BondGraphTools` [61] and the control systems library `python-control` [62]. In particular, for each example, the nonlinear ODEs are automatically generated in symbolic form from the bond graph representation. These ODEs are then automatically linearised and the corresponding transfer functions generated. The corresponding Python code for all of the examples is to be found at https://github.com/gawthrop/Oscillation24.

Although not considered here, we believe that the the analysis presented in this paper will lead to the systematic synthesis of biochemical oscillators.

## 2 Analysis of bond graph feedback

The analysis of bond graph feedback has been discussed in the context of control of biological systems [49]. This section illustrates and reformulates this approach in the context of oscillations. The basic ideas of bond graph modelling of chemical reaction networks are given elsewhere [25] and are not repeated here. In this formulation, the chemical potential *ϕ* for each species is a non-linear function of the state *x* (amount of substance) which in turn is the accumulation of the net flow *f* of the species.

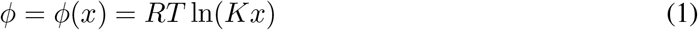

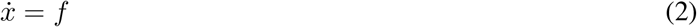

where *K* is a thermodynamic constant [24].

Figure 1(a) shown the generic bond graph feedback loop used in this paper. The component labelled **BG** represents the bond graph of an arbitrarily complex chemical reaction network which generates a product species represented by the Bond Graph component **Ce**:**P**. This product also drives the chemical reaction network, thus forming a (bond graph) feedback loop. The **0** junctions imply that:

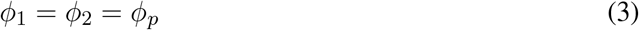

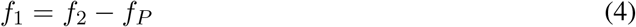

**Figure 1:**
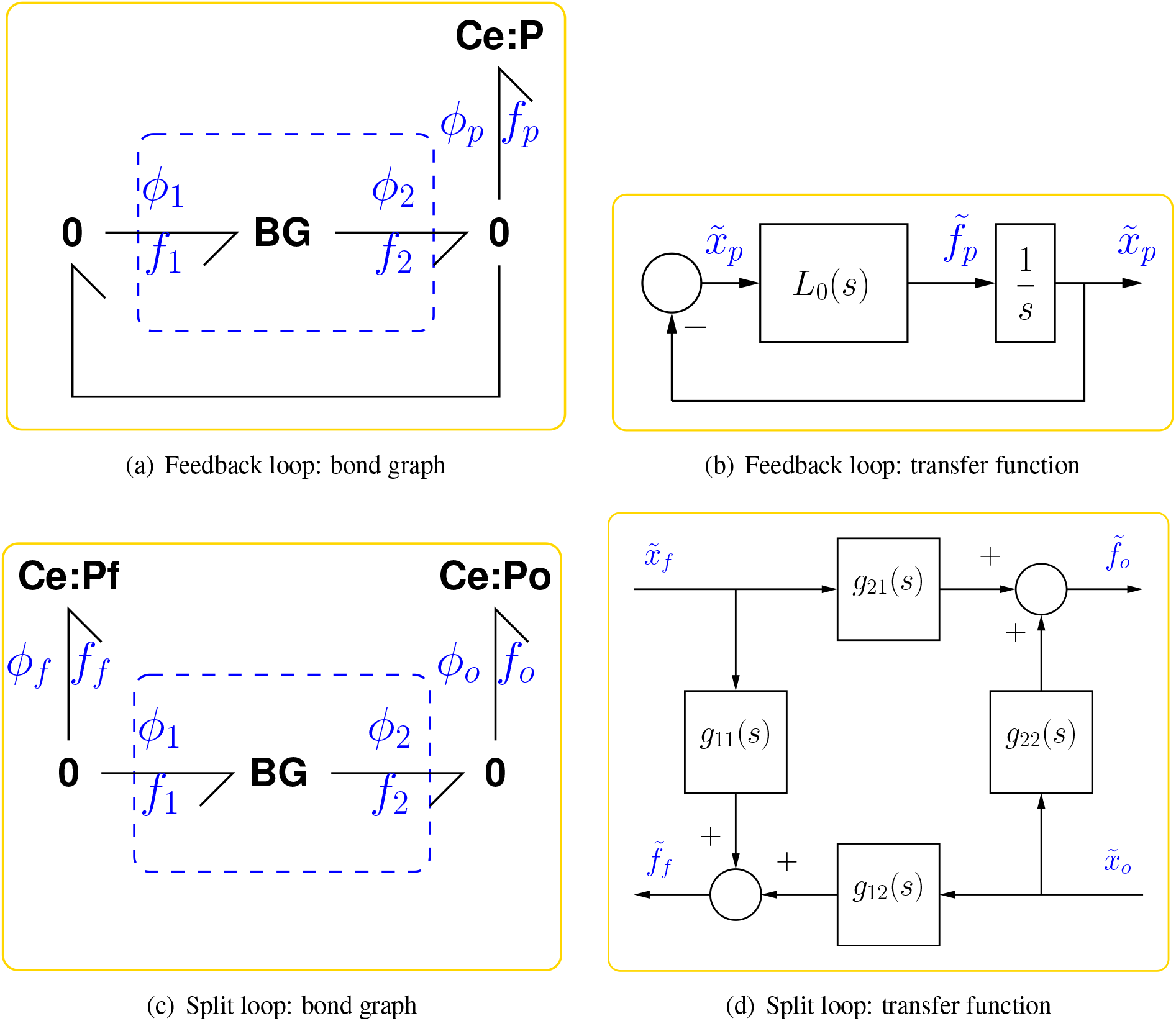
Analysis of bond graph feedback. The chemical potential *ϕ*(*x*) for each species is a non-linear function of the state *x* (amount of substance) which in turn is the accumulation of the net flow *f* of the species. (a) The basic bond graph feedback system [49] where **BG** is a (possibly complex) bond graph submodel and **Ce**:**P** represents the product. (b) The transfer function based equivalent of (a), which is used for feedback analysis and is equivalent to Equations (6) & (7). (c) The split-loop bond graph [49] used to define active and passive feedback; the two components **Ce**:**Pf** and **Ce**:**Po** have identical properties to **Ce**:**P** of (a) and the **0** junction connections imply that *f*_*f*_ = − *f*_1_ and *f*_*o*_ = *f*_2_. (d) The transfer function equivalent of (c) defined in Equation (9) and, as discussed in the text, arranged as an amplifier with output 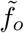 and input 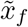. As discussed in the text, (d) and (b) are related by 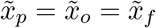 and 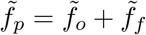 and the transfer function *L*_0_(*s*) of (b) is the negative sum of the four transfer functions in (d); −*g*_21_ is denoted the *active* component of *L*_0_(*s*).

### 2.1 Linearisation

Linearisation considers small perturbations 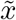 of a variable *x* about a steady-state value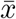, thus:

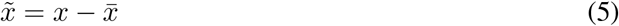

As discussed elsewhere [28] the nonlinear ODE corresponding to the CRN represented by **BG** in the configuration of Figure 1(a) can be *linearised* about a steady state. In particular, if the loop is broken by setting **Ce**:**P** to be a *chemostat* [49], the incremental product flux 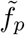 can be expressed in terms of the incremental product state 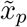 as:

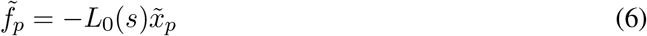

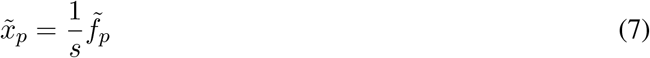

where *s* is the Laplace operator, −*L*_0_(*s*) the *transfer function* representing the CRN and the − sign is introduced for compatibility with standard negative feedback control representations. As discussed elsewhere [28], the control theory convention of linearising in terms of the state *x*, rather than the potential *ϕ*(*x*), is used. Equations (6) and (7) have the block-diagram representation of Figure 1(b).

In control theory terminology the *loop gain L*(*s*) corresponding to the feedback system of Figure 1(b) is the product of the two transfer functions:

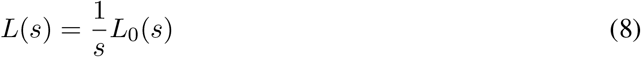

Analysis of *L*(*s*) is the basis of the frequency response analysis of control systems [47].

In this paper, the emphasis is on oscillation rather than control, however the loop-gain *L*(*s*) remains a key transfer function in the classical control systems analysis. Insight into the structure of *L*_0_ can be obtained from a *split-loop* approach [49]. In particular, with reference to Figure 1(c) an additional chemostat **Ce**:**Pf**, representing the feedback of product *P*, is added.

With reference to Figure 1(b) the conventional feedback loop carries a single variable and thus the loop gain *L*(*s*) is a transfer function with one input and one output. In contrast, the feedback bond of Figure 1(a) carries two variables and thus, when linearised, corresponds to a matrix transfer function with two inputs and two outputs and therefore with four scalar transfer functions as elements. In particular, the nonlinear ODE corresponding to the CRN represented by **BG** in the configuration of Figure 1(c) can be *linearised* about a steady state to give the four transfer functions of Figure 1(d):

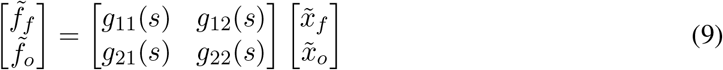

For the bond graphs of Figures 1(a) & 1(c) to be equivalent:

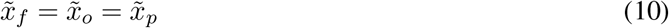

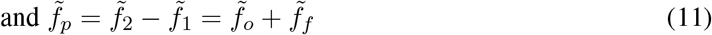

hence

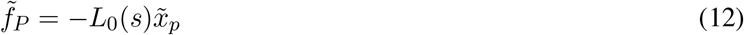

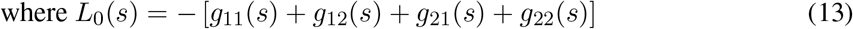

Thus *L*_0_ is the negative sum of the four transfer functions. In engineering terms, the system represented by **BG** can be thought of as an *amplifier* generating flow *f*_*o*_ from state *x*_*f*_ and the four transfer functions can be interpreted as:

*g*_21_ The forward gain

*g*_12_ The reverse gain

*g*_11_ The input admittance

*g*_22_ The output admittance

In engineering terms, an ideal amplifier would be defined by zero reverse gain, input admittance and output admittance: *g*_12_ = *g*_11_ = *g*_22_ = 0; thus the forward gain *g*_21_ can be regarded as the active part of the amplifier and other terms the passive parts. In biological terms, this ideal amplifier would correspond to a reaction network displaying no retroactivity [57–59] – which is impossible in practice; by analogy, the terms active and passive used in the biological context. For this reason, and following Gawthrop [49], the transfer function *L*_0_ is decomposed into the two transfer functions *L*_0act_ and *L*_0pas_ where:

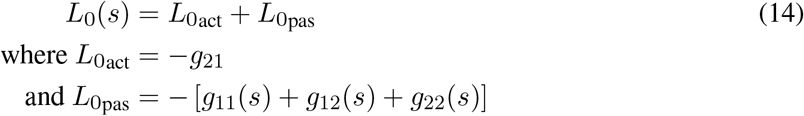

Once again, the negative sign is introduced to follow the negative feedback convention of control theory in Figure 1(b).

Similarly, the loop-gain *L* is decomposed as:

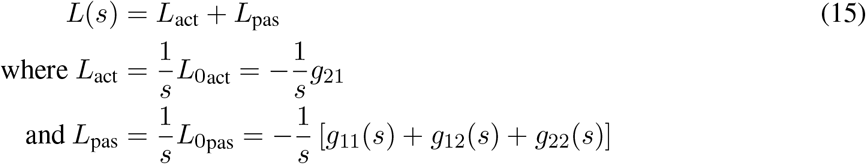

As an example, consider the reaction

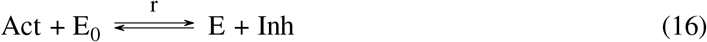

represented by the bond graph of Figure 2(d). This reaction represents enzyme activation and inhibition. Assuming mass action, the reaction flux is:

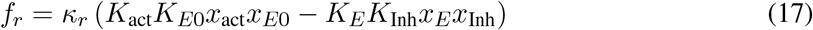

**Figure 2:**
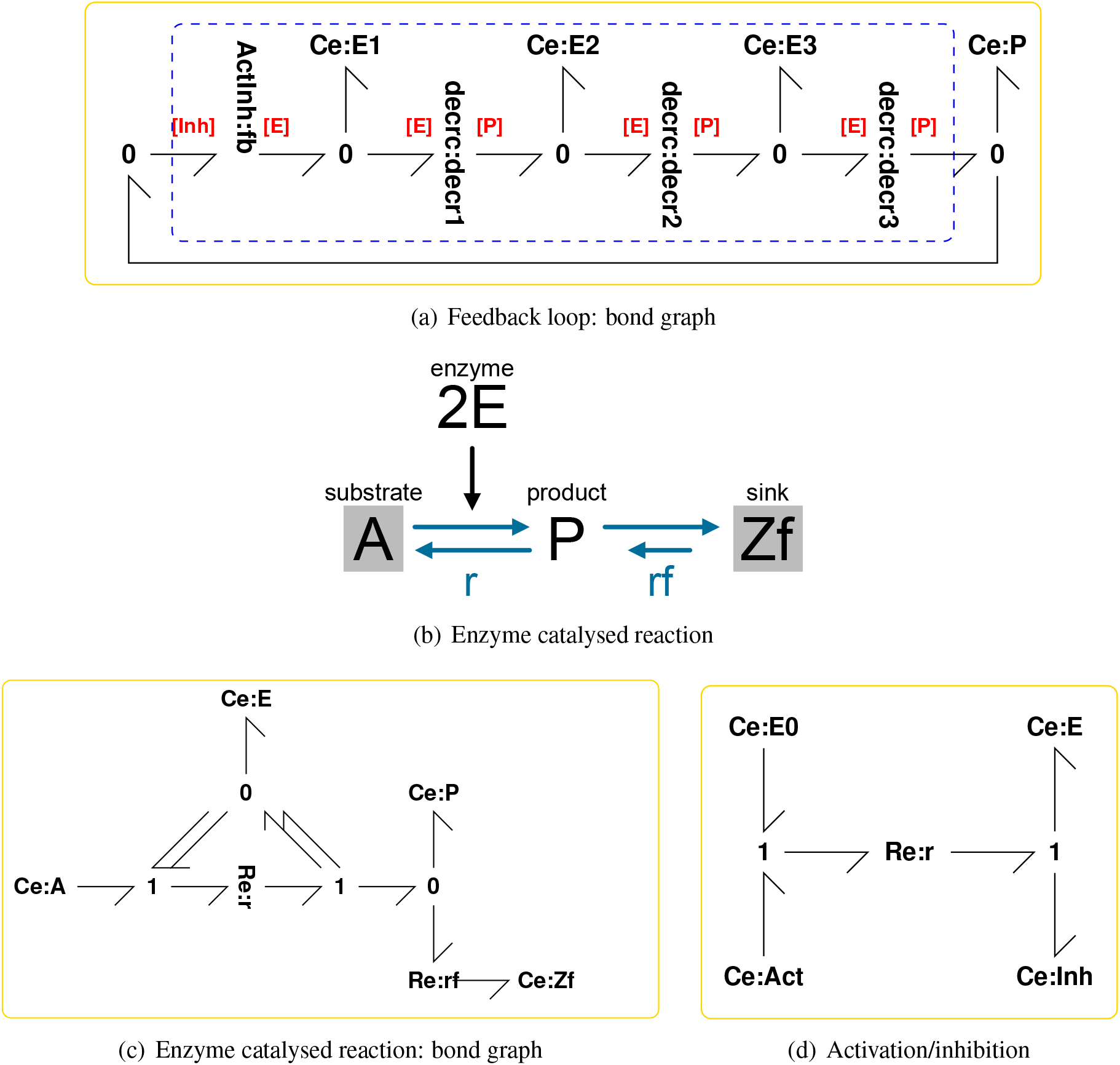
Illustrative example: system model. (a) A particular example of the generic feedback loop of Figure 1(a) comprising three instances of the bond graph submodule of Figure 2(c) representing an enzyme catalysed reaction and the bond graph submodule of Figure 2(d) representing an activation/inhibition reaction. This is a *negative* feedback loop where the product P *inhibits* the reaction. (b) The reaction diagram for each of the three enzyme catalysed reactions embedded in (a); the corresponding bond graph appears in (c). (c) The bond graph of the enzyme catalysed reaction discussed else-where [24] augmented by a degradation reaction **Re**:**rf** and with two-fold cooperativity indicated by the double bonds. **Ce**:**A, Ce**:**P** and **Ce**:**E** represent the substrate, product and enzyme respectively; **Re**:**r** represents the corresponding reaction. **Ce**:**Zf** and **Re**:**rf** represent product degradation where **Ce**:**Zf** is a chemostat with near-zero chemical potential. (d) A activation inhibition reaction where **Ce**:**E0, Ce**:**E, Ce**:**Act** and **Ce**:**Inh** represent the bound enzyme, the unbound enzyme, the activator and the inhibitor respectively.

Choosing the input port to be the inhibitor Inh and the output port to be the enzyme E, and noting that the flow *f*_*r*_ is common to each port:

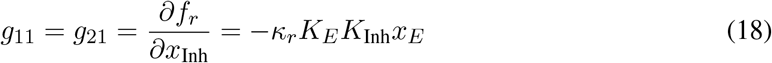

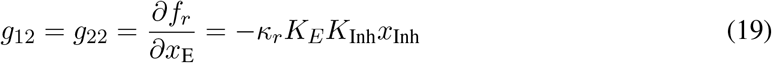

Note that these two transfer functions each depend on the steady-state values

As a further example, consider the bond graph of Figure 2(c). This represents the enzyme catalysed reaction with cooperativity and degradation:

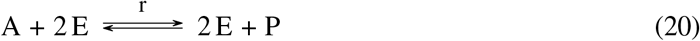

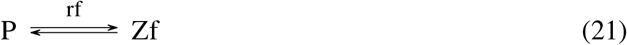

where E is the enzyme (the controlling input in this case), A the substrate and P the product (the output in this case). The reaction labelled *r* is the enzyme catalysed reaction and the reaction labelled *rf* is the degradation reaction. The species Zf represents the degradation sink with zero potential. Both **Ce**:**A** and **Ce**:**Zf** are chemostats for the purposes of this example. The corresponding reaction flows are:

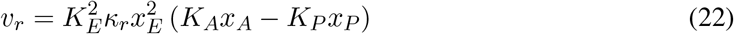

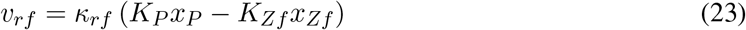

The net enzyme flow *v*_*E*_ is zero and the net product flow is *v*_*p*_ = *v*_*r*_ − *v*_*rf*_ hence:

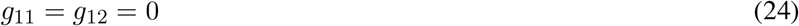

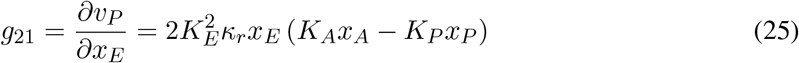

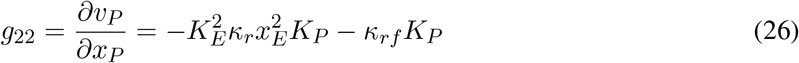

In this case, the forward gain *g*_21_ can be made large, and the output admittance *g*_22_ made relatively small by choosing a large value for *K*_*A*_*x*_*A*_. (thus rendering the reaction approximately irreversible) and a small value for *κ*_*rf*_. Furthermore, the forward gain can be increased by cooperativity and replacing the double bonds by more than two bonds.

### 2.2 Steady-state computation

Section 2.1 finds the linearisation of the nonlinear system of Figure 1(a) to give the transfer function representation of Figure 1(b). This requires a steady-state of the system where all species amounts *x* (states) are constant.

In general, it is difficult to find an unstable steady-state of a nonlinear system. In this paper, it is assumed that the nonlinear system represented by **BG** of Figure 1(a) is stable and thus the transfer function *L*_0_(*s*) of Figure 1(b) is stable; thus instability is assumed to be induced by the feedback. The approach taken here is to set the product (P) component **Ce**:**P** to be a chemostat thus making the corresponding state *x*_*P*_ a constant. In general, the corresponding flow *f*_*P*_ ≠ 0 thus closing the loop would not yield a steady-state. The value of *x*_*P*_ that gives *f*_*P*_ = 0 in the steady-state is found by iteratively solving simulating the open-loop system to a steady-state and using a standard root-finding algorithm `fsolve` within the Python package `scipy.optimize`. The simulation takes account of conserved moieties as discussed in § 3c of [24].

Details are given in the code at https://github.com/gawthrop/Oscillation24.

### 2.3 Frequency response, phase margin and stability

As discussed in § 2.1, the linearised system is represented by the loop-gain transfer function *L*(*s*) where *s* is the Laplace operator. It is standard control engineering practice [47] to replace *s* by *jω* where 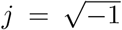 is the unit imaginary number and *ω* is the frequency with units of rad s^−1^. At each frequency *ω, L*(*jω*) is a complex number with gain (magnitude) *L*(*jω*) and phase (in degrees) ∠*L*(*jω*)°.

This frequency response can be used to examine the stability properties of a system via the Bode Diagram [47]. This approach is simplified when the loop gain *L*(*s*) corresponds to a stable system; and this is the case for each of the systems considered in this paper.

The Bode diagram of *L*(*s*) consists of two logarithmic plots against frequency *ω*: gain |*L*(*jω*)| and phase ∠*L*(*jω*). For example, Figures 3(a) & 3(b) show the two parts of the Bode diagram for the three transfer functions: *L*(*s*) (Total), *L*_act_ (Active) and *L*_pas_ (Passive). The critical frequency *ω*_*c*_ is the value of *ω* where |*L*(*jω*)| = 1; this is marked on Figure 3(a). The corresponding phase margin *θ*_*pm*_ is

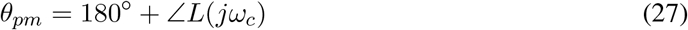

**Figure 3:**
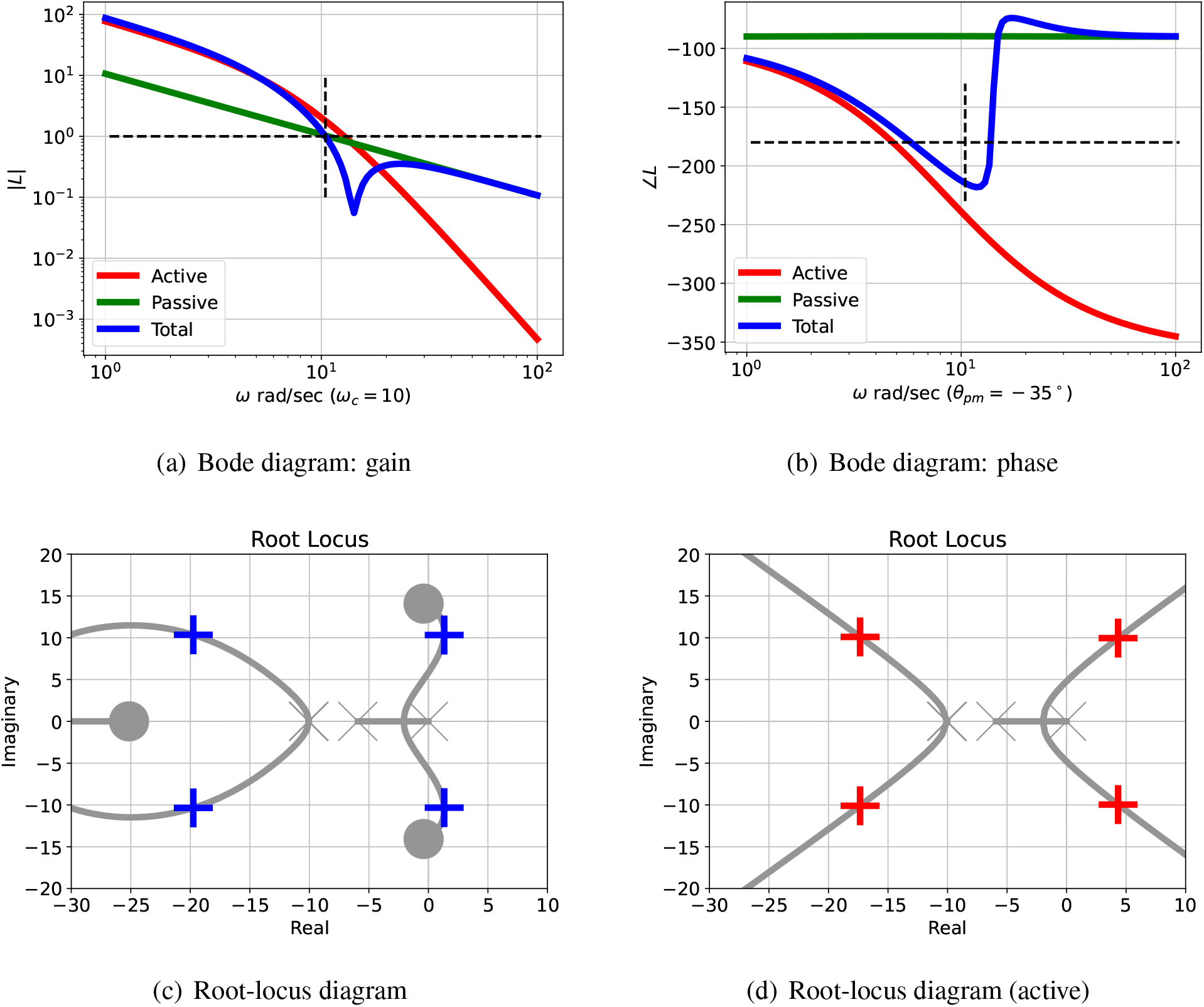
Illustrative example.: analysis. (a) The Bode magnitude plot of the transfer function gains (on a logarithmic scale) against frequency for three cases: |*L*_act_(*jω*)|, |*L*_pas_(*jω*)| and |*L*(*jω*)|. The frequency *ω*_*pm*_ where the gain |*L*(*jω*) = 1 is marked on the diagram. (b) The corresponding Bode phase plot of ∠*L*_act_(*jω*), ∠*L*_pas_(*jω*) and ∠*L*(*jω*). The phase margin of *L*(*jω*) is ∠*L*(*jω*_*pm*_) + 180° In this case, the phase margin is *θ*_*pm*_ = − 35° at *ω*_*pm*_ = 10.44 rad s^−1^; the negative sign indicating instability. (c) The Root Locus diagram corresponding to *L*(*s*). The *open-loop* poles and zeros are marked as × and◯ respectively; the *closed-loop* poles are marked as **+** or **+** (active only). Poles with positive real parts correspond to exponentially increasing responses and complex poles correspond to oscillatory responses. (d) The Root Locus diagram corresponding to *L*_act_(*s*).

This is marked on Figure 3(b). A negative phase margin indicates instability and thus the possibility of oscillation.

Other stability measures are available, including the gain margin and the stability margin [47, § 9.3]; these could be used as an alternative to the phase margin.

## 3 Illustrative example

The stability properties of linearised system can be analysed by any of the multitude of methods drawn from control theory [47] based on the feedback system loop-gain transfer function *L*(*s*). This paper focuses on two such methods: the Bode diagram and the root-locus diagram. The Bode diagram is based on the frequency response *L*(*jω*) of *L*(*s*) and thus can be used for system of arbitrarily high order; the root locus provides further insights but is restricted to low-order systems or, as discussed in § 5, a reduced order system.

These ideas are illustrated by the example of this section; the Glycolytic Oscillator of Sel’kov [11] is discussed in § 4 and the Repressilator of Tyson et al. [63], as modelled by Pan et al. [16] is discussed in § 5.

### 3.1 System model

This illustrative example is similar to that of Goodwin [10] and has the bond graph representation of Figure 2 which is a particular example of the generic feedback loop of Figure 1(a). The state equations were automatically derived from the bond graph and are listed, together with the parameters, in the Supplementary Material S2.

This example illustrates the following points:

- Active and passive feedback components of the feedback loop where:
  – The active component gives rise to an unstable, oscillatory linearised system
  – The passive component tends to stabilise the system; therefore if the ratio of passive to active component is too large, the oscillation will be quenched.
- Oscillation is dependent on system *parameters* which affect the ratio of passive to active feedback.
- Oscillation is dependent on system *structure* such as number of reactions and degree of cooperativity which affect the ratio of passive to active feedback.

### 3.2 Linear analysis

The parameter values are

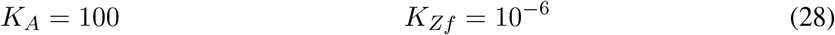

with all other parameter values unity; the small value of *K*_*Zf*_ ensures that the chemostat represented by **Ce**:**Zf**n near-zero chemical potential. The corresponding system steady state values were all unity except for:

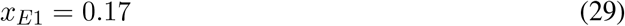

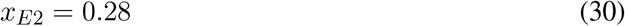

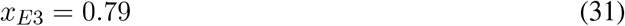

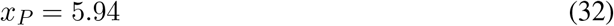

Applying the numerical linearisation to give the transfer functions of Equation (9) as:

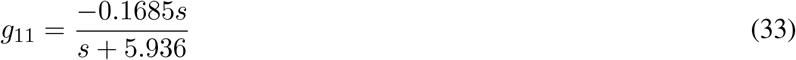

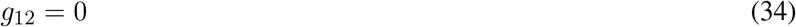

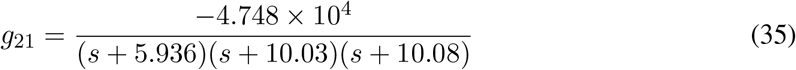

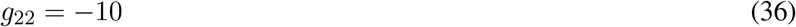

Note that *g*_11_ and *g*_21_ contain the Laplace operator *s* and thus represent dynamical systems of first and third order respectively.

Hence, using equation (15):

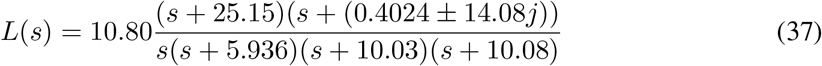

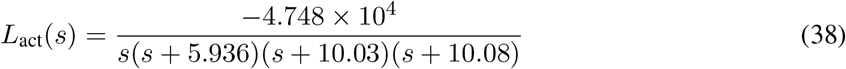

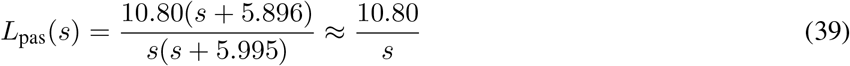

Figure 3(a) shows the Bode magnitude plot for the active (38) and passive (39) components of the loop gains together with the total loop gain (37). Figure 3(b) shows the corresponding Bode phase diagram.

The passive portion of the loop gain, *L*_pas_, has little effect on the overall transfer function at low frequencies (including this value of *ω*_*pm*_), but dominates at higher frequencies. The phase margin *θ*_*pm*_ is thus highly dependent on the properties of *L*_pas_ and thus system parameters such as *κ*_*rf*_ appearing in Equation (26). In this case, the phase margin *θ*_*pm*_ = − 35 ° at a frequency of *ω*_*pm*_ = 10.44 rad s^−1^; the negative phase margin indicates instability. The dependency of the phase margin *θ*_*pm*_ on *κ*_*rf*_ is shown in Figure 5(a) for a three values of *K*_act_.

The root-locus diagram corresponding to *L*_act_ appears in Figure 3(d) and the root-locus diagram corresponding to *L* appears in Figure 3(c); the closed-loop poles are marked by the red crosses. One pair of closed loop poles is complex (indicating an oscillatory system) and is in the the right-half plane (indicating instability) at *s* = 1.157 *±* 10.37*j*. As in the Bode analysis, the passive portion of the loop gain, *L*_pas_, has little effect on the closed-loop poles in this case. The effect of adding *L*_pas_ (39) to *L*_act_ (38) is to give a loop gain *L* (37) with zeros at *s* = − 25.15 and *s* = − (0.4024 *±* 14.08*j*). The corresponding three branches of the root locus diagram terminate at these zeros; whereas *L*_act_ (38) has no finite zeros and the corresponding branches of the root locus diagram tend to infinity. This linear analysis indicates that the system steady state is unstable with an oscillatory response.

Figure 4(a) shows the linear and nonlinear responses when the steady-state is perturbed by a small amount. As expected, the initial responses are close, but diverge as time increases. A larger initial perturbation (not shown) would lead to divergence starting at a smaller time. Figure 4(b) shows the nonlinear response when the steady-state is perturbed plotted as a phase-plane diagram. As expected, the initial response is a spiral corresponding to the linear oscillation, but the the response settles to a limit cycle; the linear response (not shown) spirals outwards without bound.

**Figure 4:**
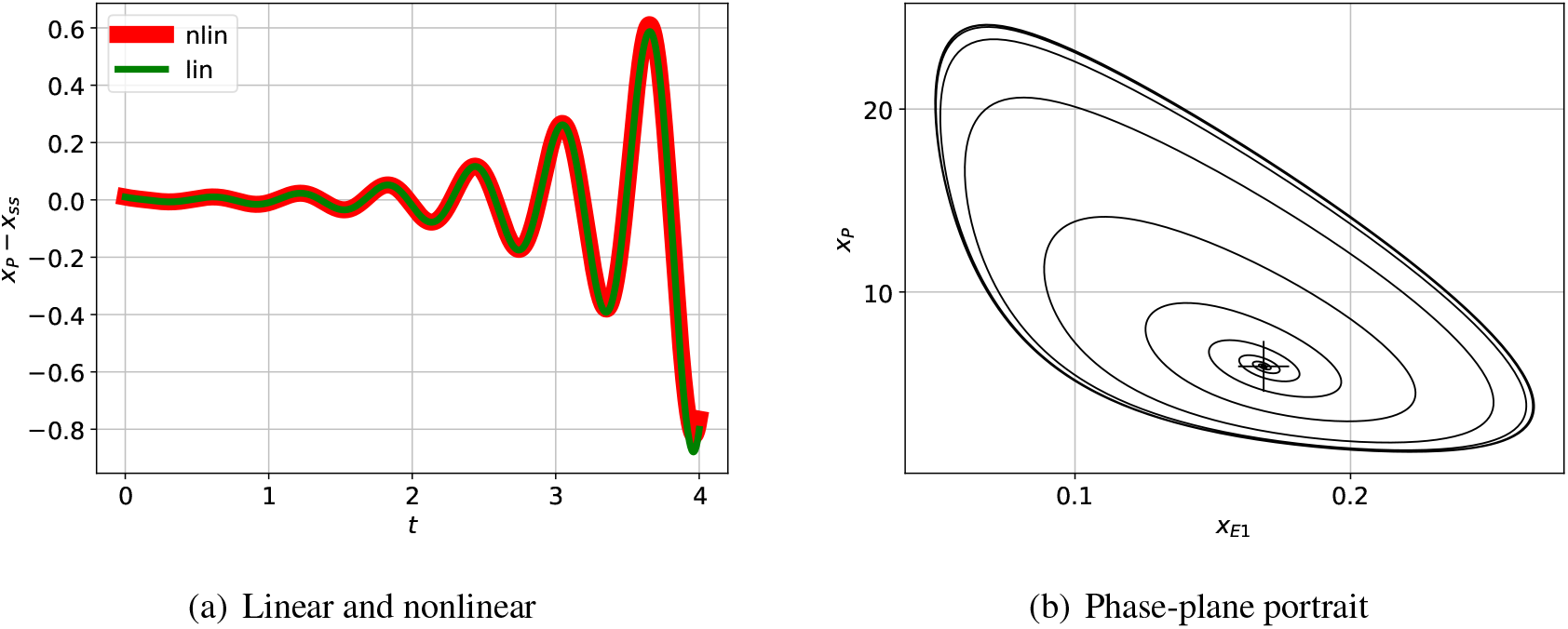
Illustrative example.: nonlinear simulation. (a) The incremental response of the indicated state to an initial condition corresponding to the perturbed steady-state and plotted against time *t* for a short time span. As expected, the initial nonlinear and linear responses are close for the small initial perturbation; as indicated in (b) the reponses diverge as time increases. (b) The phase-plane response corresponding to two particular states plotted for a longer time. The steady-state value is marked by +. The trajectory spirals outwards from the initial condition to the limit cycle.

### 3.3 Interplay of passive and active feedback

The oscillation illustrated in Figures 3 & 4 is dependent on the value of the system parameters. Figure 5(a) shows how the system phase margin *θ*_*pm*_ of the linearised system depends on two parameters: *K*_act_ and *κ*_*rf*_. The range of these parameters where *θ*_*pm*_ ≥ 0 corresponds to a stable system with no possibility of oscillation. Figures 5(a) – 5(d) give the phase margin *θ*_*pm*_ as parameters vary for three systems with modified structure.

**Figure 5:**
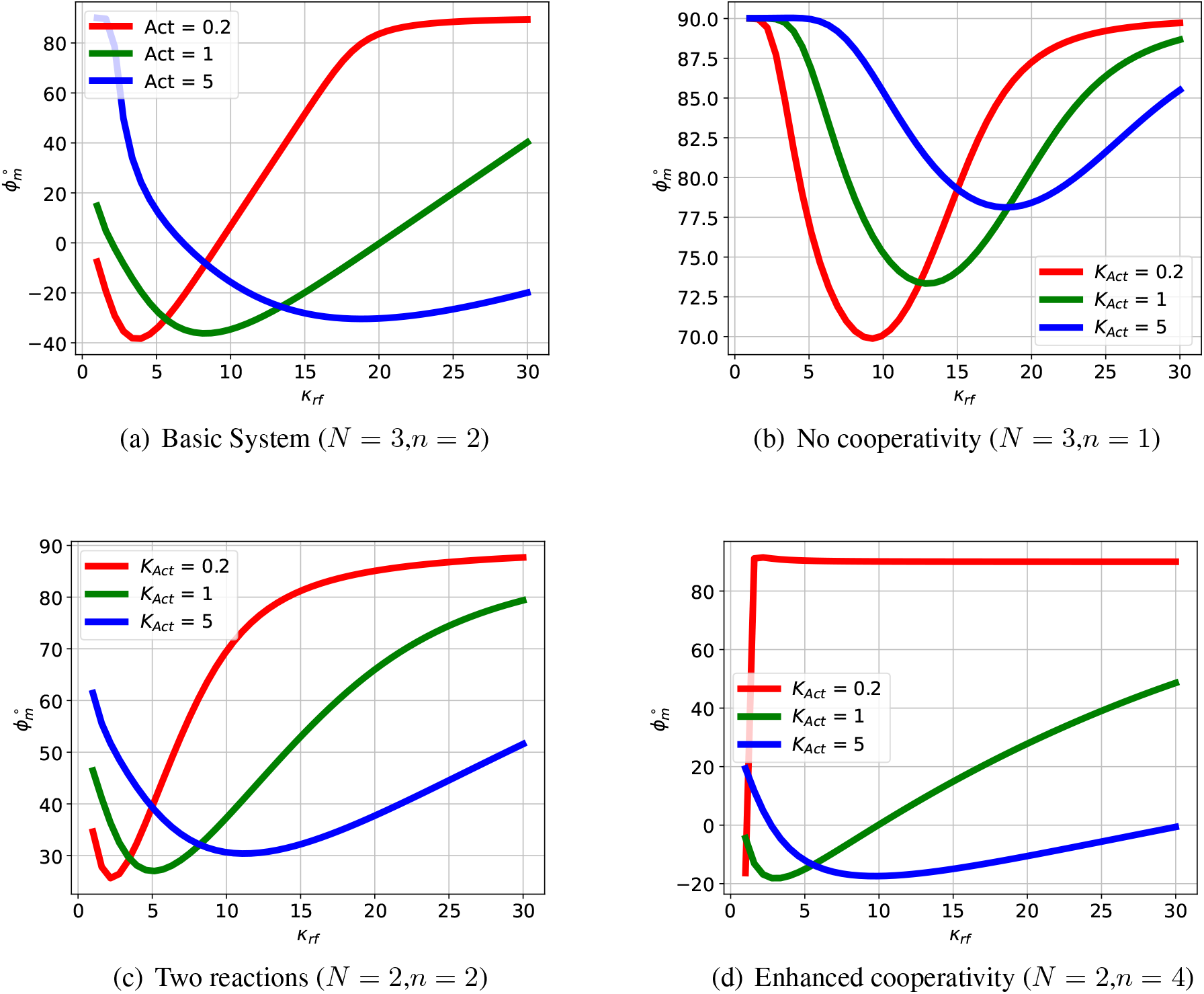
Illustrative Example: phase margin. The phase margin *θ*_*pm*_ is a stability indicator derived from the frequency response of the feedback loop; a positive value indicating stability and therefore no oscillation. (a) The basic system with three reactions (*N* = 3) and cooperativity (*n* = 2) has a range of parameters with negative phase margin *θ*_*pm*_ giving potential oscillation. For each value of *K*_act_ (which drives the reactions), there is a different value of the degradation parameter *κ*_*rf*_ giving a maximum negative value of *θ*_*pm*_. (b) If the basic system is modified to have no cooperativity (*n* = 1), the phase margin *θ*_*pm*_ is positive for all values of *K*_act_ and *κ*_*rf*_ so there is no oscillation. (c) If the basic system is modified to have two enzyme-catalysed reactions (*N* = 2) the phase margin *θ*_*pm*_ is positive for all values of *K*_act_ and *κ*_*rf*_ so there is no oscillation. (d) If the basic system is modified to to have two enzyme-catalysed reactions (*N* = 2) and enhanced cooperativity (*n* = 4) there is a range of parameters with negative phase margin *θ*_*pm*_ giving potential oscillation. As in (a), for each value of *K*_act_ (which drives the reactions), there is a different value of the degradation parameter *κ*_*rf*_ giving a maximum negative value of *θ*_*pm*_.

As can be seen from these figures, increasing the number *N* of enzyme-catalysed reactions enhances the possibility of oscillations; this is because the *phase-lag* − ∠*L*_act_ of the active feedback *L*_act_ increases with *N*. Increasing the number *n* of cooperativity bonds enhances the possibility of oscillations; this is because the *gain* |*L*_act_| of the active feedback *L*_act_ increases with *n*.

The variation of *κ*_*rf*_ and *K*_act_, as indicated in Figure S1 (Supplementary Material), affects the system steady-state and, because the system is nonlinear, this affects the linearised system as well. In this case small and large values of *κ*_*rf*_ correspond to no oscillation, and the range is modulated by the value of *K*_act_.

As indicated in Figures 3(a) & 3(b), the phase margin *θ*_*pm*_, and thus possibility of oscillation, depends on the interplay of the frequency response of the active and passive feedback. Therefore a closer understanding of the conditions for oscillation relies on how both system parameters and system structure alter the relative contribution of the active and passive feedback.

As indicated in Figure 5(a), the closed loop system is stable when *K*_act_ = 0.2 and *κ*_*rf*_ = 10. This case is examined in more detail in Figures 6(a) & 6(b) and compared to the the case where *K*_act_ = 1 and *κ*_*rf*_ = 10. In this particular case, although the the active part of the loop-gain described by *L*_act_ corresponds to an unstable closed-loop system, its magnitude is decreased so that the unchanged magnitude of the the passive part *L*_pas_ is large enough for the overall loop-gain described by *L* to give a stable closed-loop system. Thus, in this case, the passive component has the unwanted effect of quenching the oscillation.

**Figure 6:**
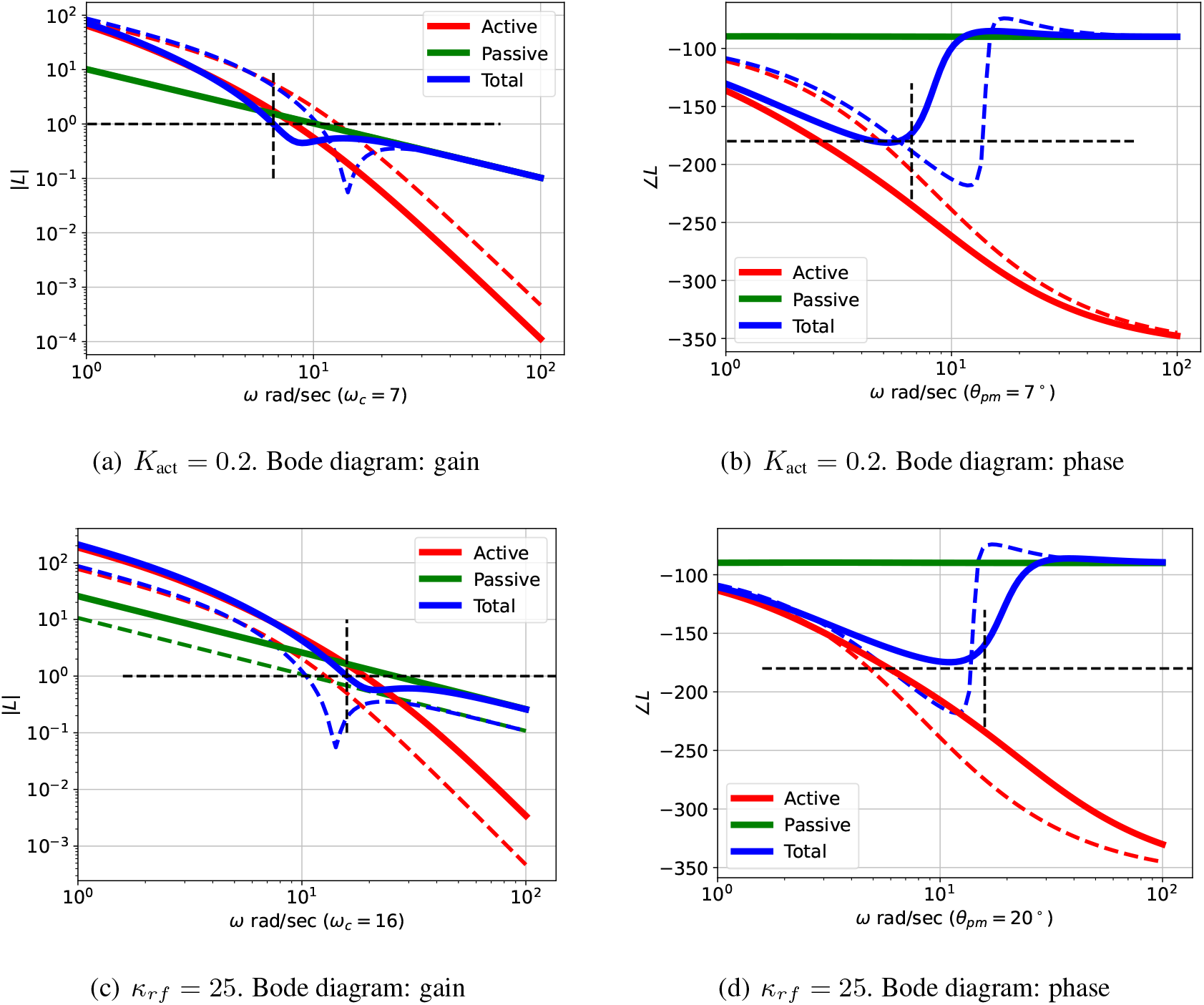
Illustrative example: Comparative Bode diagrams for parametric variation. The dashed lines correspond to the example of Figure 3, the firm lines correspond to the modified parameters given in the captions. (a) & (b) The passive part of the feedback is unchanged, but the gain of the active part is reduced so that the passive part dominates the overall feed back at the reduced critical frequency *ω*_*c*_ = 7 rad s^−1^ giving a positive phase margin (*θ*_*pm*_ = 7°) indicating no oscillation. (c) & (d) The gain of both the active and passive parts of the feedback are increased in such a way that the passive part dominates at the critical frequency *ω*_*c*_ = 16 rad s^−1^ giving a positive phase margin (*θ*_*pm*_ = 20°) indicating no oscillation.

As indicated in Figure 5(a), the closed loop system is stable when *K*_act_ = 1 and *κ*_*rf*_ = 25. This case is examined in more detail in Figures 6(c) & 6(d) and compared to the the case where *K*_act_ = 1 and *κ*_*rf*_ = 10. In this particular case, although the the active part of the loop-gain described by *L*_act_ corresponds to an unstable closed-loop system, its magnitude, the magnitude of the the passive part *L*_pas_ is increased enough to outweigh the increase in the gain of the active part. Thus, once again, the passive component has the unwanted effect of quenching the oscillation.

The interplay of passive and active feedback is dependent on system structure as well as system parameters; this is examined in Figure 7. Figures 7(a) & 7(b) show that removing cooperativity reduces the active feedback gain so that the passive feedback gives a positive phase margin thus quenching the oscillation. Figures 7(c) & 7(d) show that reducing the number of reaction stages from *N* = 3 to *N* = 2 reduces the active feedback phase so that that the passive feedback gives a positive phase margin thus quenching the oscillation.

**Figure 7:**
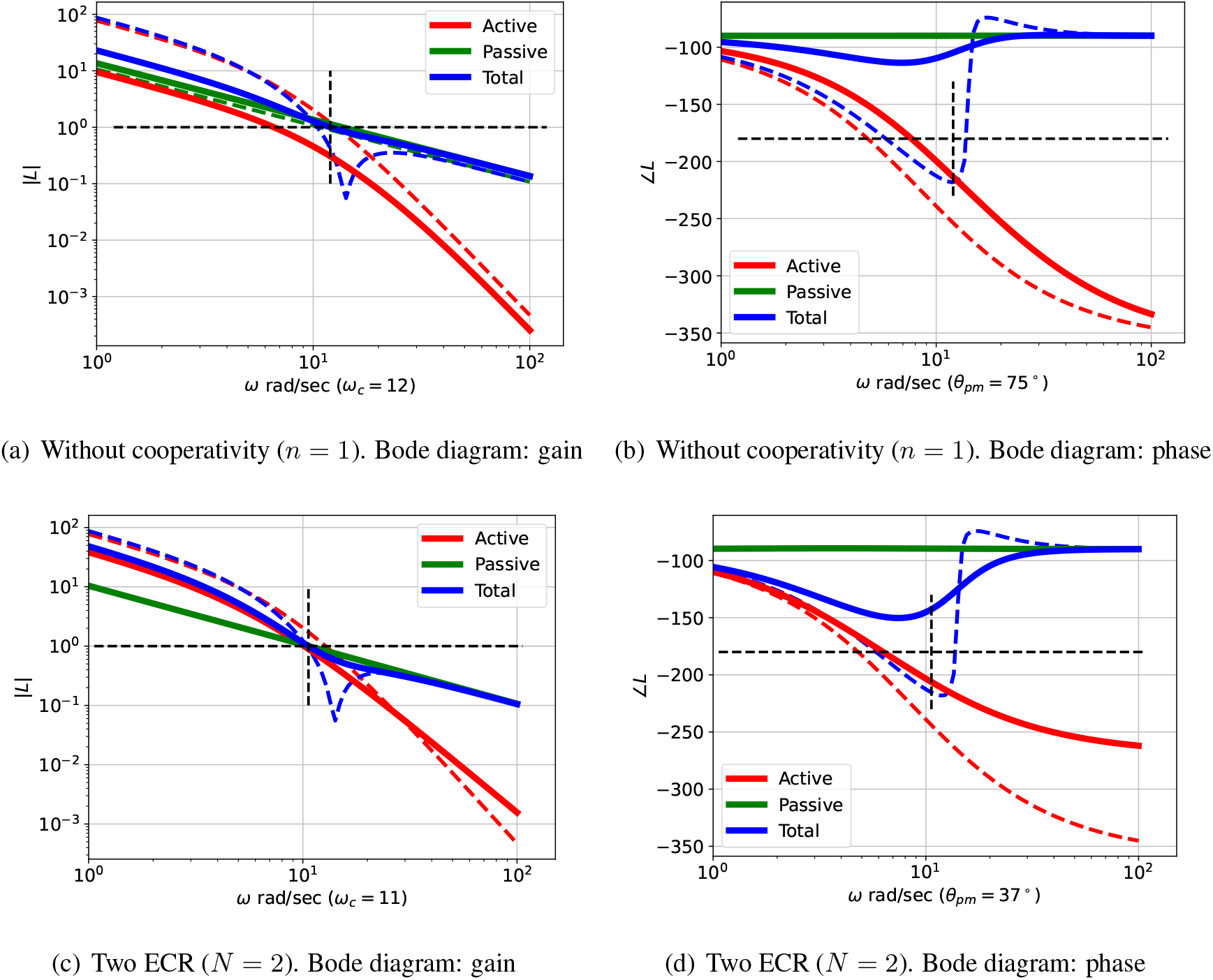
Illustrative example: Comparative Bode diagrams for structural variation. The dashed lines correspond to the example of Figure 3, the firm lines correspond to the modified structures given in the captions. (a) & (b) Without cooperativity (*n* = 1). The passive part of the feedback is largely unchanged, but the gain of the active part is reduced so that the passive part dominates the overall feed back at the reduced critical frequency *ω*_*c*_ = 12 rad s^−1^ giving a positive phase margin of *θ*_*pm*_ = 75°. (c) & (d) Two enzyme-catalysed reactions (ECR) (*N* = 2). The phase lag of the active part of the feedback is reduced in such a way that the overall phase lag at the slightly modified critical frequency *ω*_*c*_ = 11 rad s^−1^ gives a positive phase margin of *θ*_*pm*_ = 37°.

## 4 The Sel’kov Oscillator

### 4.1 System model

The Glycolytic Oscillator of Sel’kov [11] is analysed by Keener and Sneyd [12] using normalised parameters. As discussed elsewhere [24, 25], the corresponding chemical reaction network can be transformed into the bond graph of Figure 8 where Figure 8(a) is a particular example of the generic feedback loop of Figure 1(a). The double bond of Figure 8(a) corresponds to cooperativity *n* = 2 corresponding to the value *γ* = 2 given by Keener and Sneyd [12, Fig. 1.7]. The state equations were automatically derived from the bond graph and are listed, together with the parameters, in the Supplementary Material S2.

**Figure 8:**
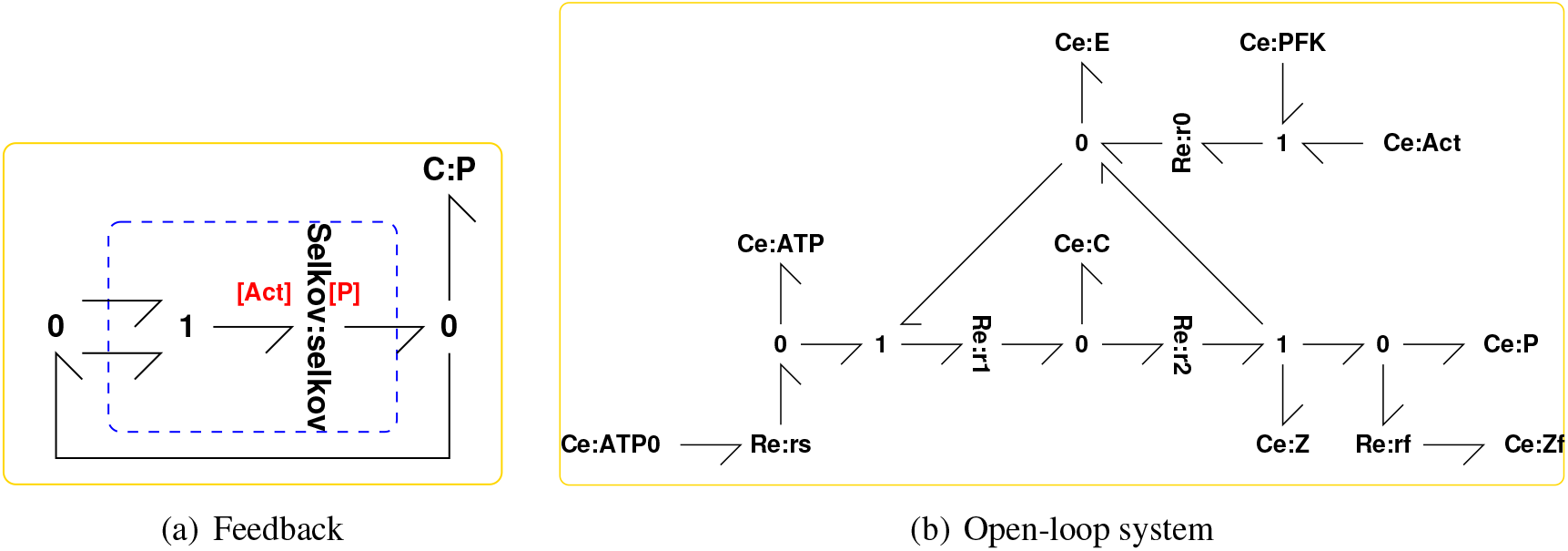
Sel’kov Glycolytic Oscillator: system model. (a) A particular example of the generic feedback loop of Figure 1(a) comprising the bond graph of (b) embedded in a *positive* feedback loop where the product P *activates* the reaction. (b) The bond graph corresponding to the the Glycolytic Oscillator Sel’kov [11] as presented by Keener and Sneyd [12]. The flow of ATP is determined by the chemostat **Ce**:**ATP0** and the reaction **Re**:**rs** with the parameters of Table 1. The chemostat **Ce**:**Z** with near-zero potential makes the reaction corresponding to **Re**:**r2** approximately irreversible and, as in Figure 2, the **Ce**:**Zf** and **Re**:**rf** components represent degradation.

This example illustrates:

**Table 1:** Sel’kov Glycolytic Oscillator: Normalised Parameters. The small values of *K*_*Z*_ and *K*_*Zf*_ give the corresponding chemostats near-zero potential. The normalised flow of ATP is approximately *κ*_*rs*_*K*_*AT P* 0_ = 0.6.

| Parameter | Value |
| --- | --- |
| $K_Z = K_{Zf}$ | $10^{-10}$ |
| $\kappa_{r0} = \kappa_{r1} = \kappa_{r2}$ | $10^3$ |
| $\kappa_{rf}$ | 10 |
| $K_{ATP0}$ | $10^3$ |
| $\kappa_{rs}$ | $6 \times 10^{-4}$ |

- Active and passive feedback components of the feedback loop where:
  – The active component corresponds to *positive* feedback and gives rise to an unstable, non-oscillatory linearised system
  – The passive component corresponds to *negative* feedback which would, by itself lead to a stable, but oscillatory linear system.
  – The two components together give an unstable, oscillatory system if their ratio is not too large or too small.
- As discussed in § 3.1, oscillation is dependent on both system *parameters* and system *structure*.

### 4.2 Linear analysis

In a similar fashion to the illustrative example of § 3, the transfer functions of Equation (9) describing the linearised system are:

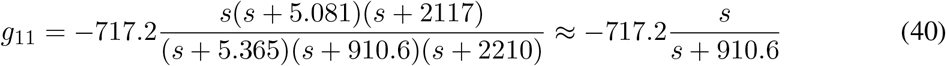

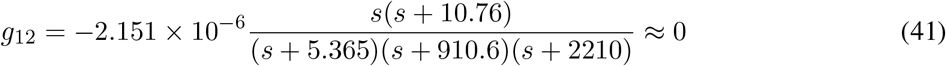

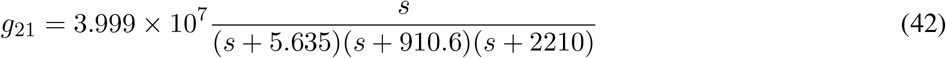

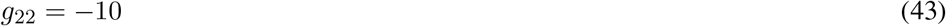

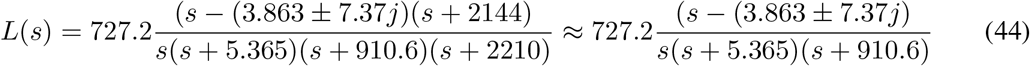

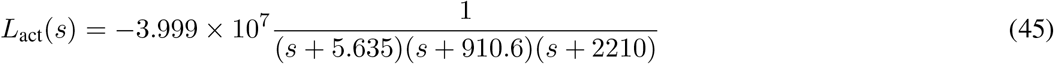

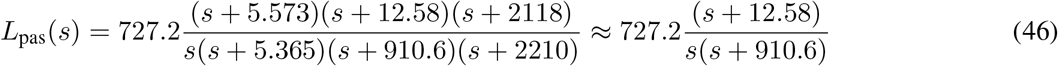

Note that *L*_act_(*s*) is preceded by a *negative* sign indicating *positive* feedback.

Figure 9(a) shows the Bode magnitude plot for the active (45) and passive (46) components of the loop gains together with the total loop gain (44). Figure 9(b) shows the corresponding Bode phase diagram.

**Figure 9:**
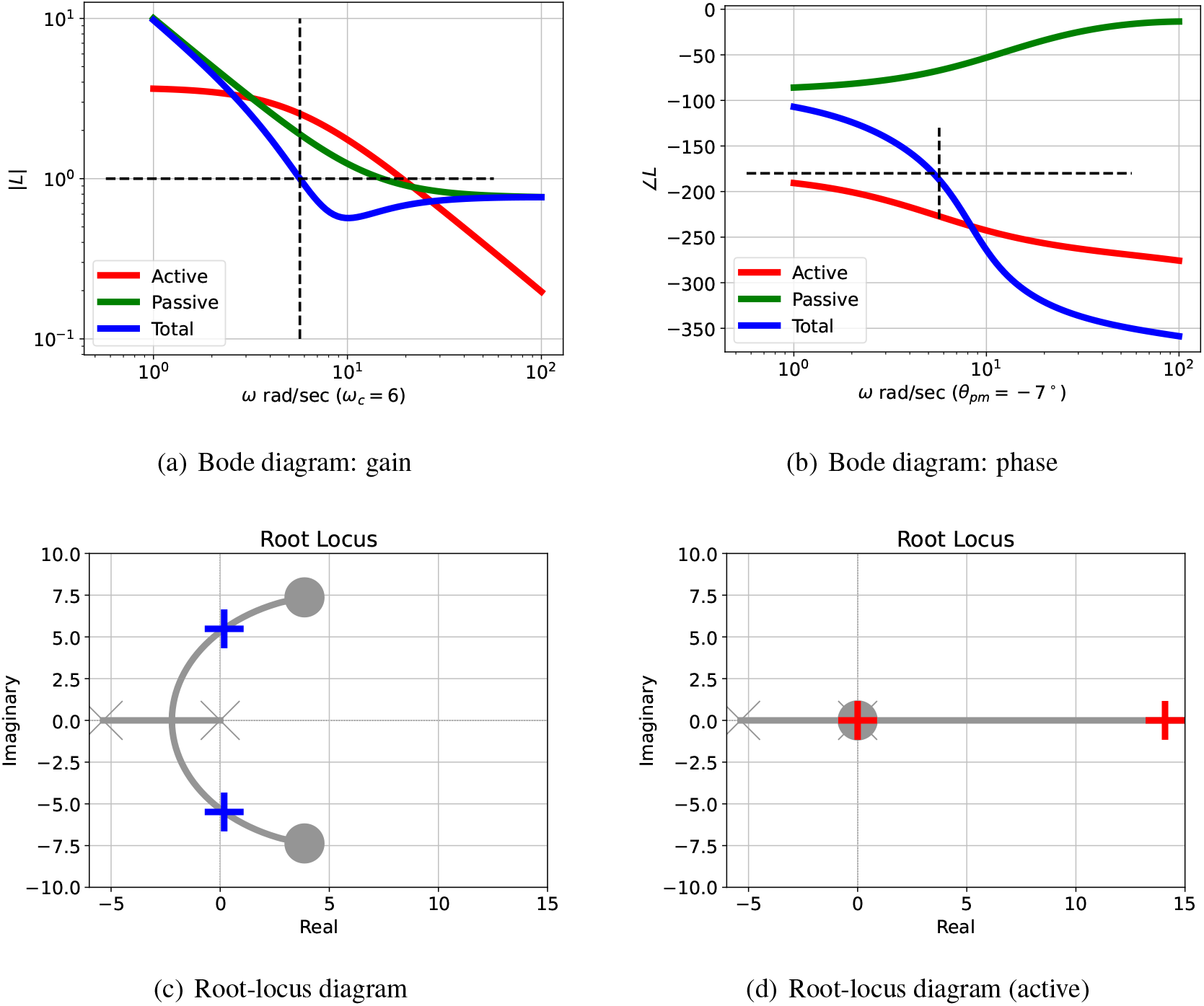
Sel’kov Glycolytic Oscillator: analysis. (a) The Bode magnitude plot of the transfer function gains (on a logarithmic scale) against frequency for three cases: |*L*_act_(*jω*)|, |*L*_pas_(*jω*)| and |*L*(*jω*)|. The frequency *ω*_*pm*_ where the gain |*L*(*jω*) = 1 is marked on the diagram. (b) The corresponding Bode phase plot of ∠*L*_act_(*jω*), ∠*L*_pas_(*jω*) and ∠*L*(*jω*). The phase margin of *L*(*jω*) is ∠*L*(*jω*_*pm*_) + 180° In this case, the phase margin is *θ*_*pm*_ = − 7° at *ω*_*pm*_ = 5.69 rad s^−1^; the negative sign indicating instability. (c) The Root Locus diagram corresponding to *L*(*s*). The *open-loop* poles and zeros are marked as × and ◯ respectively; the *closed-loop* poles are marked as **+** or **+** (active only). Poles with positive real parts correspond to exponentially increasing responses and complex poles correspond to oscillatory responses. (d) The Root Locus diagram corresponding to *L*_act_(*s*).

Figure 10(a) shows the linear and nonlinear responses when the steady-state is perturbed. As expected, the initial responses are close, but diverge as time increased.

**Figure 10:**
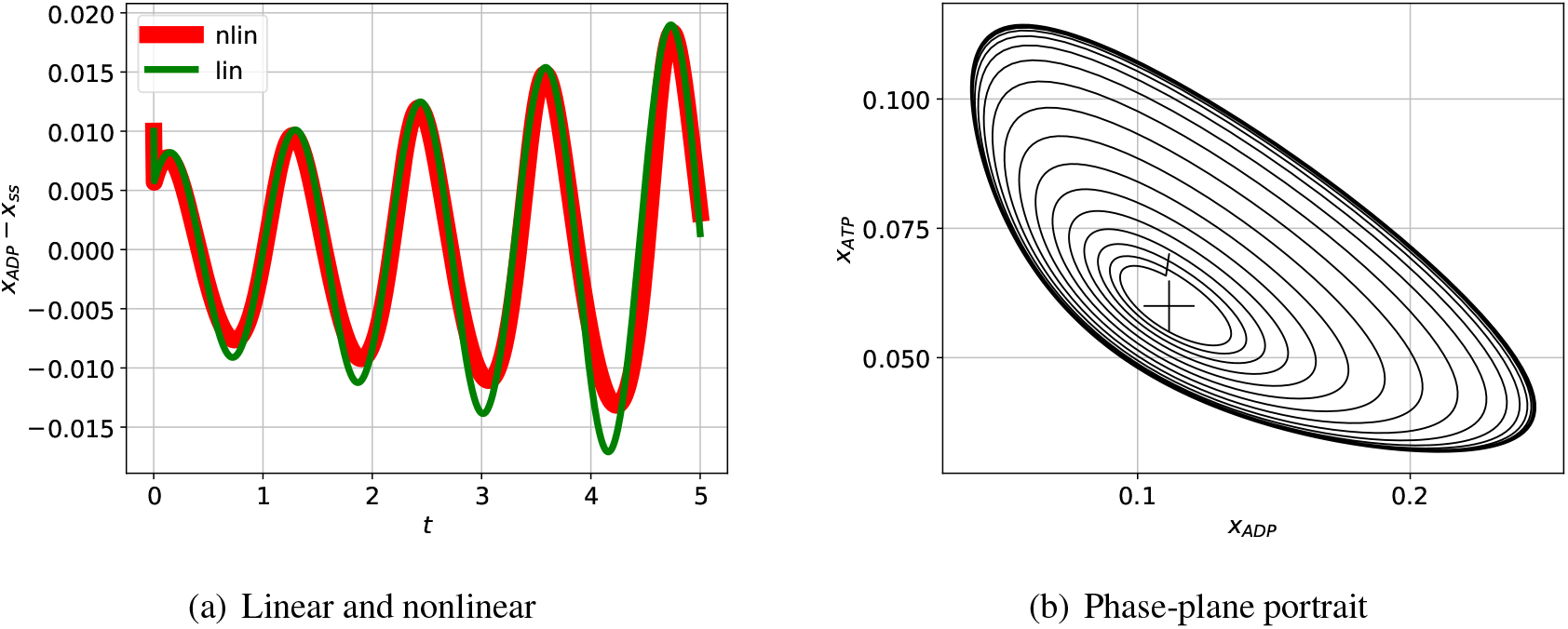
Sel’kov Glycolytic Oscillator: nonlinear simulation. (a) The incremental response of the indicated state to an initial condition corresponding to the perturbed steady-state and plotted against time *t* for a short time span. As expected, the initial nonlinear and linear responses are close for the small initial perturbation; as indicated in (b) the reponses diverge as time increases. (b) The phase-plane response corresponding to two particular states plotted for a longer time. The steady-state value is marked by +. The trajectory spirals outwards from the initial condition to the limit cycle.

### 4.3 Interplay of passive and active feedback

The passive portion of the loop gain, *L*_pas_, dominates at both low and high frequencies. However, unlike the illustrative example of § 3, the critical frequency *ω*_*pm*_ occurs in the mid-frequency region where neither *L*_pas_ nor *L*_act_ dominates – it is this interaction of positive feedback (from *L*_act_) and negative feedback (from *L*_pas_ that leads to the oscillatory response.

The root-locus diagram corresponding to *L*_act_ appears in Figure 9(d) and the root-locus diagram corresponding to *L* appears in Figure 9(c); the closed-loop poles are marked by the red crosses. In the case of Figure 9(c) one pair of closed loop poles is complex (indicating an oscillatory system) and is in the the right-half plane (indicating instability) at *s* = 1.831 *±* 5.485*j*. Because the net loop-gain has zeros with positive real parts, it is a non-minimum phase system which leads to instability at high feedback gains [47]; these zeros arise from the addition of the positive and negative feedback paths.

However, in the case of Figure 9(d), there is a single real closed-loop pole in the left-half plane indicated non-oscillatory instability without the effect of the passive portion of the feedback loop. Unlike the illustrative example of § 3, the passive portion of the loop gain, *L*_pas_, has a significant effect on the closed-loop poles: the *positive* feedback from *L*_act_ combined with the *negative* feedback from *L*_pas_ gives rise to the oscillation.

In a similar fashion to § 3.3, Figure 5, Figure 11 shows how the phase margin *θ*_*pm*_ varies with two parameters, in this case *v*_*AT P*_, the flow of ATP and *κ*_*rf*_, the degradation parameter. The basic system with cooperativity (*n* = 2) has a range flows *v*_*AT P*_ which give rise to a negative phase margin *θ*_*pm*_ giving potential oscillation for each of the three values of *κ*_*rf*_. However for no cooperativity (*n* = 1) *θ*_*pm*_ *>* 0; the feedback system is stable and no oscillation is possible. On the other hand, increased cooperativity (*n* = 3) causes the positive feedback to dominate giving *θ*_*pm*_ *<* 0 and thus instability for all parameter values; as discussed in Figure 9, this gives rise to a single positive real pole giving instability without oscillation.

**Figure 11:**
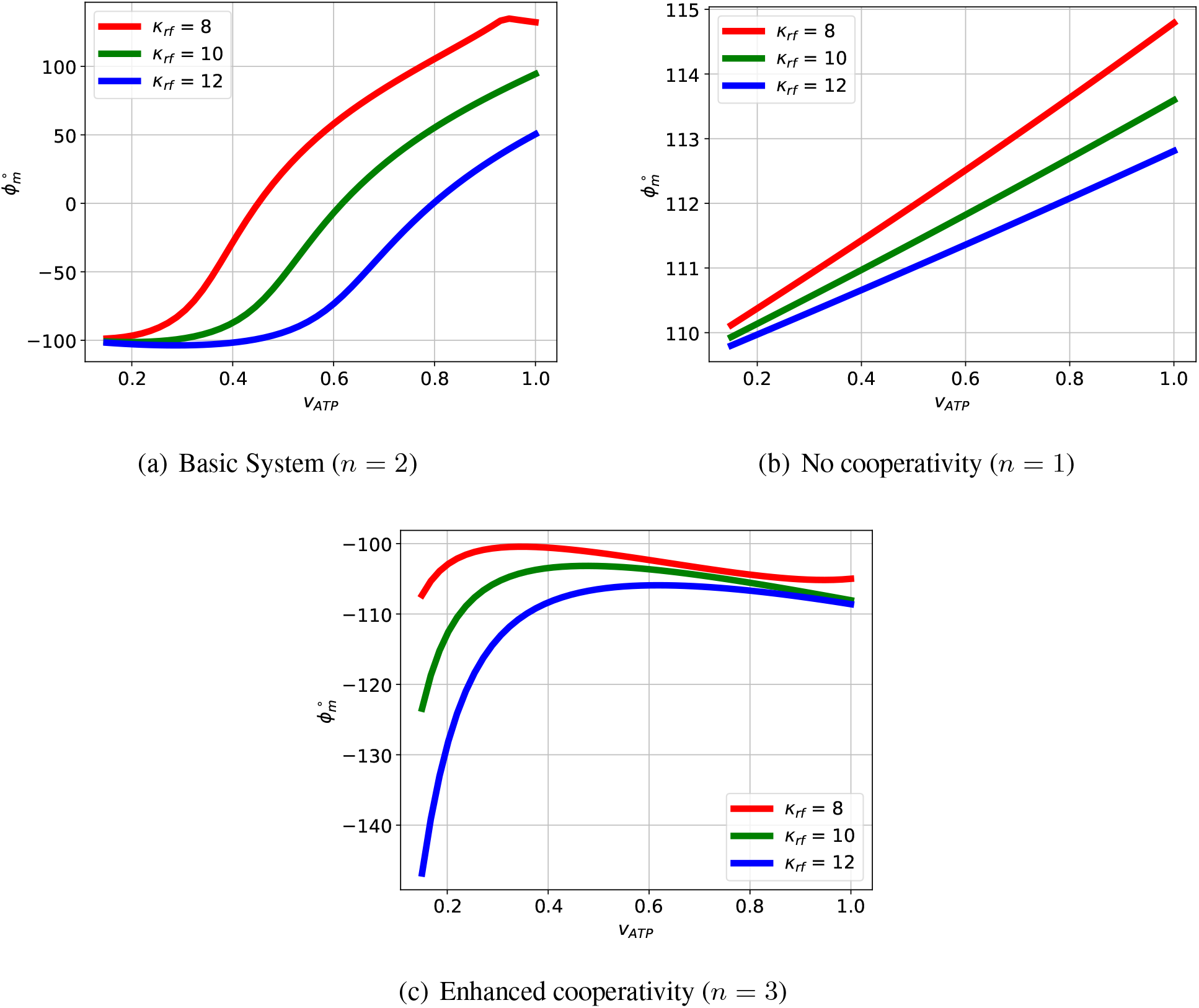
Sel’kov Glycolytic Oscillator: phase margin *θ*_*pm*_. (a) The basic system with cooperativity (*n* = 2) has a range of parameters with negative phase margin *θ*_*pm*_ giving potential oscillation. For each value of the degradation parameter *κ*_*rf*_ there is a different value of the flow *v*_*AT P*_ giving a maximum negative value of *θ*_*pm*_. (b) If the basic system is modified to have no cooperativity (*n* = 1), the phase margin *θ*_*pm*_ is positive for all values of *v*_*AT P*_ and *κ*_*rf*_ so there is no oscillation. (c) If the basic system is modified to give enhanced cooperativity (*n* = 3), the phase margin *θ*_*pm*_ is negative for all values of *v*_*AT P*_ and *κ*_*rf*_ however, as discussed in the text, the pole with positive real part is real, and so there is no oscillation.

In a similar fashion to § 3.3, Figures 6 & 7, Figures 12 & 13 give comparative Bode diagrams for parametric and structural variation respectively. These diagrams emphasise that oscillation is only possible if the structure and parameters are such that the *positive* feedback from *L*_act_ and the *negative* feedback from *L*_pas_ have similar magnitudes around the critical frequency *ω*_*c*_.

**Figure 12:**
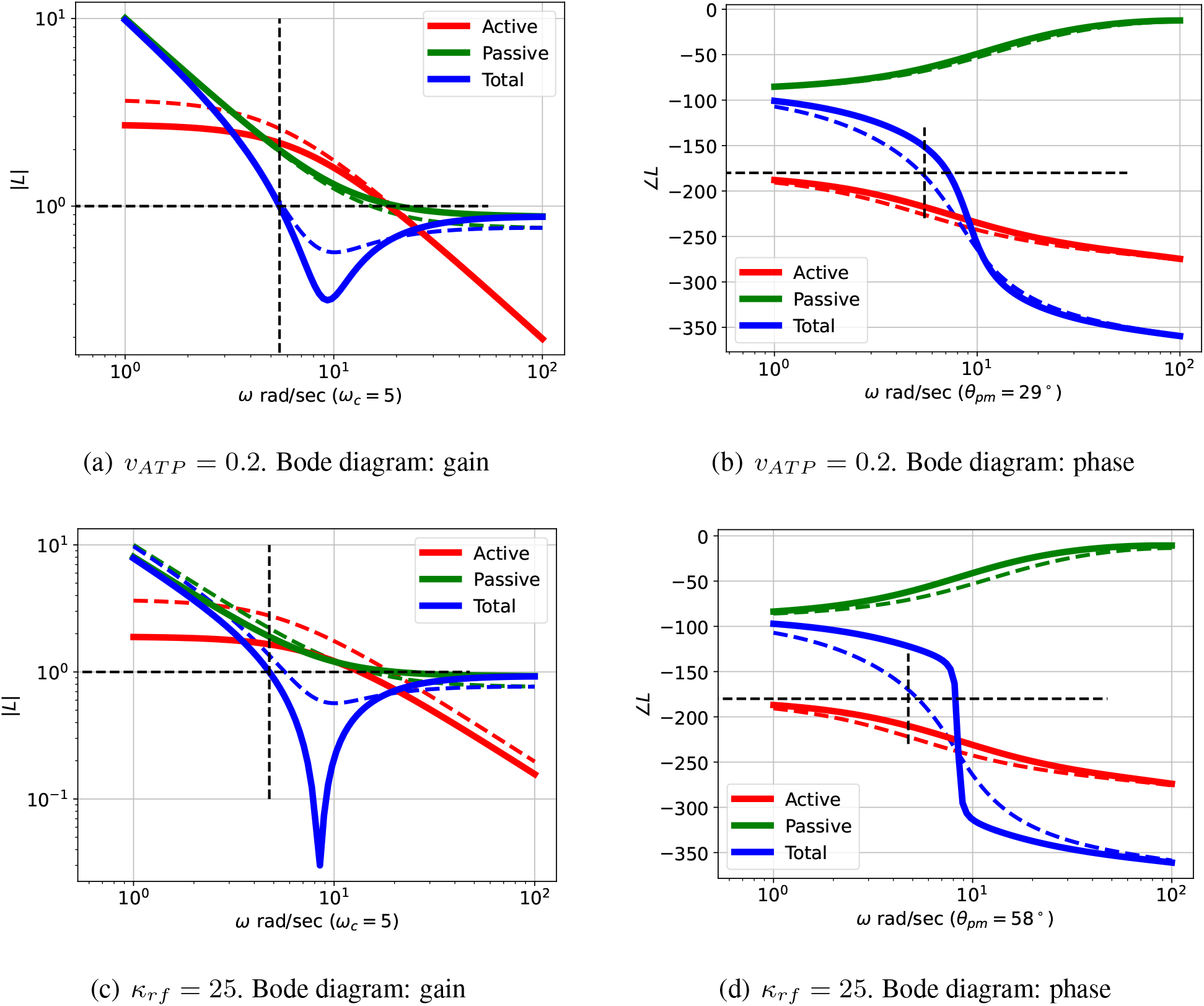
Sel’kov Glycolytic Oscillator: Comparative Bode diagrams for parametric variation. The dashed lines correspond to the example of Figure 9, the firm lines correspond to the modified parameters given in the captions. (a) & (b) The passive part of the feedback is unchanged, but the gain of the active part is reduced so that the passive part dominates the overall feed back at the critical frequency *ω*_*c*_ = 5 rad s^−1^ giving a positive phase margin (*θ*_*pm*_ = 29°) indicating no oscillation. (c) & (d) The passive part of the feedback is unchanged, but the gain of the active part is reduced so that the passive part dominates the overall feedback at the critical frequency *ω*_*c*_ = 5 rad s^−1^ giving a positive phase margin (*θ*_*pm*_ = 58°) indicating no oscillation.

**Figure 13:**
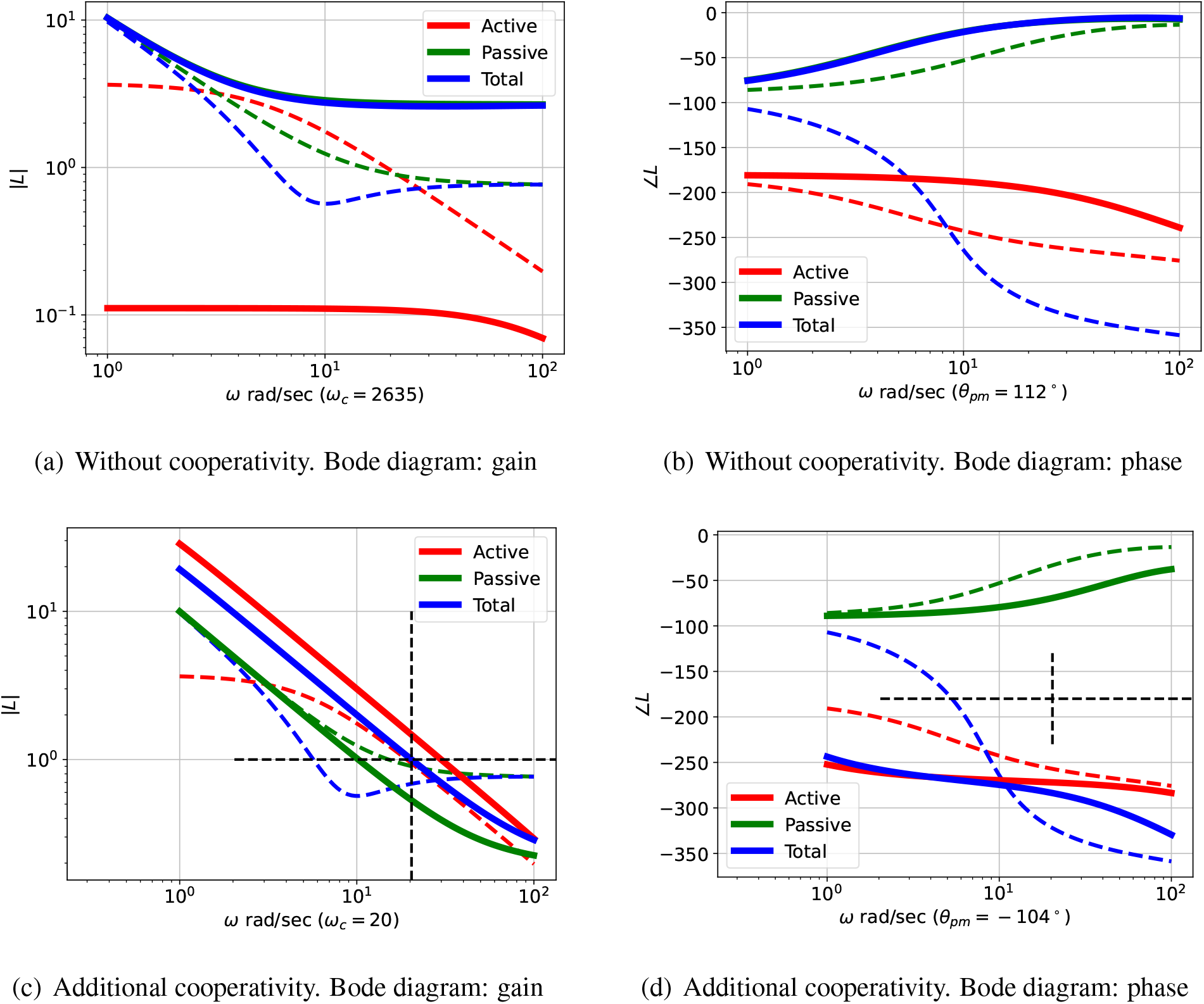
Sel’kov Glycolytic Oscillator: Comparative Bode diagrams for structural variation. The dashed lines correspond to the example of Figure 9, the firm lines correspond to the modified structures given in the captions. (a) & (b) The reduced gain of the active feedback part means that the overall feedback is dominated by the passive part. The critical frequency *ω*_*c*_ = 2365 rad s^−1^ is outside the range given and corresponds to a positive phase margin (*θ*_*pm*_ = 112°) indicating no oscillation. (c) & (d) The increased gain of the active feedback part means that the overall feedback is dominated by the active part. The critical frequency *ω*_*c*_ = 20 rad s^−1^ corresponds to a negative phase margin of (*θ*_*pm*_ = − 104°) indicating instability. However, as indicated in Figure 9(d), the active feedback corresponds to a closed-loop system with real poles, so there is no oscillation in this case; a simulation appears in Figure S3 (Supplementary Material).

## 5 The Repressilator

### 5.1 System model

A synthetic oscillator using DNA transcription, known as the repressilator, was introduced by Tyson et al. [14] and a bond graph model of the repressilator is given by Pan et al. [16]. The repressilator is a complex system and was simplified for the purposes of this example by giving the mRNAs a length of 8 (2 codons, with 4 ATP molecules per codon): the corresponding bond graph model [16] has 53 species, of which 7 are chemostats, giving 46 states. However, as shown below, the linearised system of equation (9), and thus *L*_0_ (14), can be well-approximated by a third order system. This reduction is achieved in two stages: pole/zero cancelling pairs are removed using the minimal realisation `minreal` function and the order reduced to three using the balanced order reduction `balred` function of the control systems library `python-control` [62].

The parameters from [16] are included in the Supplementary Material S2. The state equations were automatically derived from the bond graph and are also listed in the Supplementary Material S2.

This example illustrates:

- Application of the approach to a high-order system model.
- The fact that a reduced-order linearised system can be used for analysis.
- Otherwise, the system behaves similarly to the Illustrative Example of § 3.

### 5.2 Linear analysis

In a similar fashion to the illustrative example of § 4, the transfer functions of Equation (9) describing the *reduced-order* linearised system are:

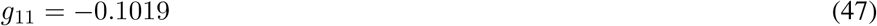

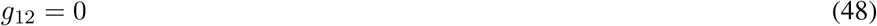

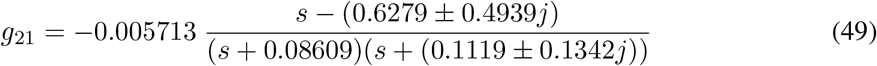

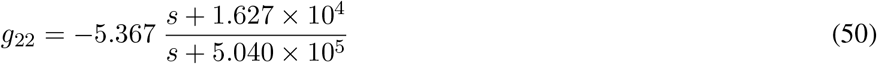

Hence, using equation (15):

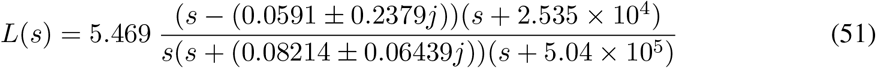

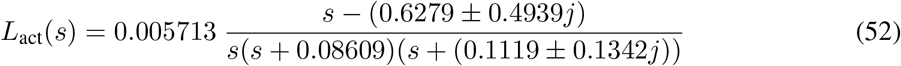

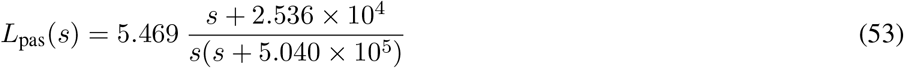

### 5.3 Interplay of passive and active feedback

Figure 14(a) shows the Bode magnitude plot for the active (52) and passive (53) components of the loop gains together with the total loop gain (51). Figure 14(b) shows the corresponding Bode phase diagram.

**Figure 14:**
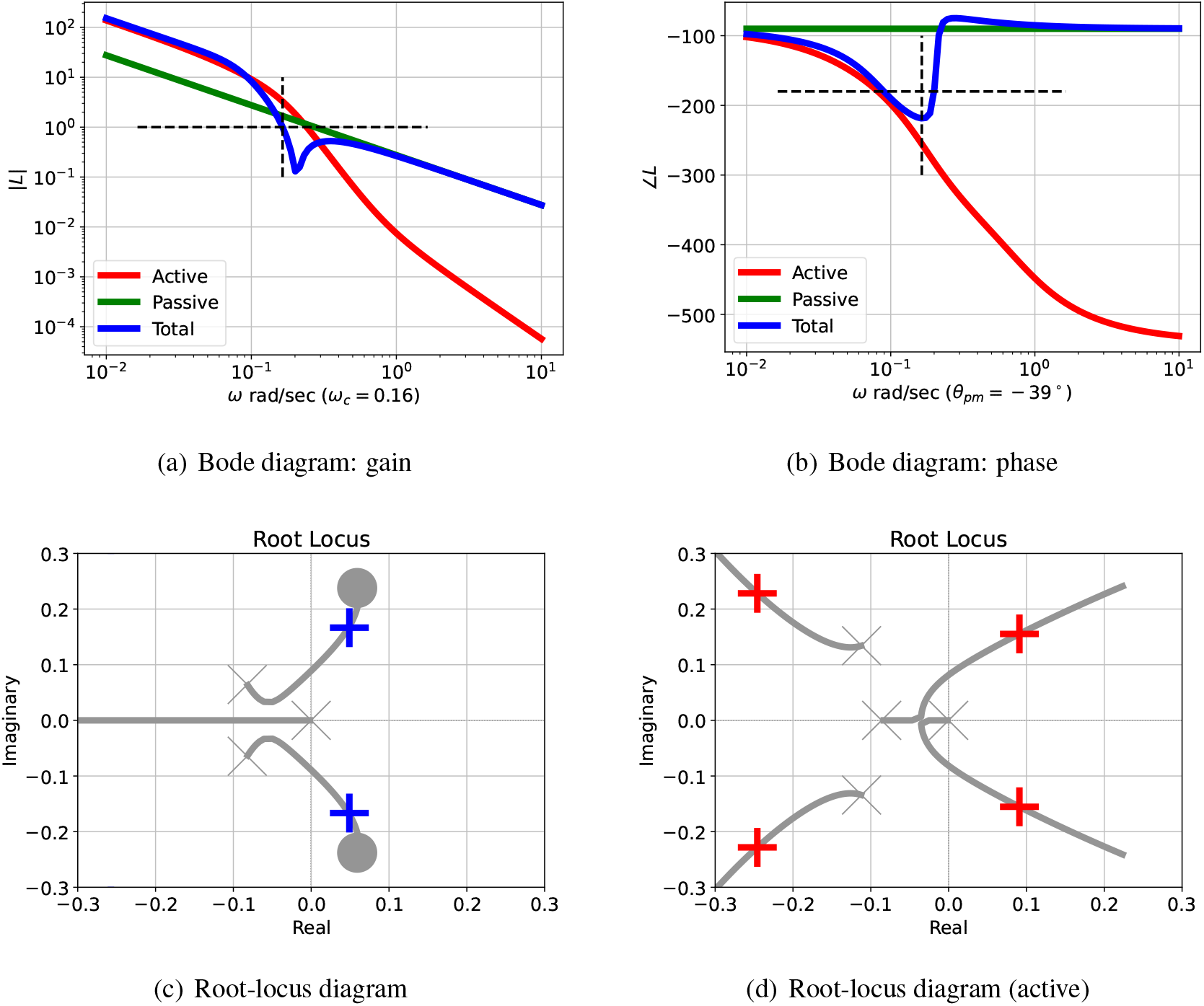
Repressilator: analysis. (a) The Bode magnitude plot of the transfer function gains (on a logarithmic scale) against frequency for three cases: |*L*_act_(*jω*)|, |*L*_pas_(*jω*)| and |*L*(*jω*)|. The frequency *ω*_*pm*_ where the gain |*L*(*jω*) = 1 is marked on the diagram. (b) The corresponding Bode phase plot of ∠*L*_act_(*jω*), ∠*L*_pas_(*jω*) and ∠*L*(*jω*). The phase margin of *L*(*jω*) is ∠*L*(*jω*_*pm*_) + 180° In this case, the phase margin is *θ*_*pm*_ = − 39° at *ω*_*c*_ = 0.16 rad s^−1^; the negative sign indicating instability. (c) The Root Locus diagram corresponding to *L*(*s*). The *open-loop* poles and zeros are marked as × and ◯ respectively; the *closed-loop* poles are marked as **+** or **+** (active only). Poles with positive real parts correspond to exponentially increasing responses and complex poles correspond to oscillatory responses. (d) The Root Locus diagram corresponding to *L*_act_(*s*).

The passive portion of the loop gain, *L*_pas_, has little effect on the overall transfer function at low frequencies (including this value of *ω*_*c*_), but dominates at higher frequencies. In this case, the phase margin *θ*_*pm*_ = − 39° at a frequency of *ω*_*c*_ = 0.16 rad s^−1^.; the negative phase margin indicates instability.

The root-locus diagram corresponding to the *reduced-order L*_act_ appears in Figure 14(d) and the root-locus diagram corresponding to the *reduced-order L* appears in Figure 14(c); the closed-loop poles are marked by the red crosses. One pair of closed loop poles is complex (indicating an oscillatory system) and is in the the right-half plane (indicating instability) at *s* = 0.04879 *±* 0.1685*j*. As in the Bode analysis, the passive portion of the loop gain, *L*_pas_, has little effect on the closed-loop poles in this case. The effect of adding *L*_pas_ (53) to *L*_act_ (52) is to give a loop gain *L* (51) with zeros at *s* = − 2.535 × 10^4^ and *s* = (0.0591 *±* 0.2379*j*). The corresponding three branches of the root locus diagram terminate at these zeros; whereas *L*_act_ (52) has no finite zeros and the corresponding branches of the root locus diagram tend to infinity. This linear analysis indicates that the system steady state is unstable with an oscillatory response. Figure 15(a) shows the linear and nonlinear responses when the steady-state is perturbed. As expected, the initial responses are close, but diverge as time increased. The response of the linear reduced model is plotted as a dashed line; despite the order reduction, the response is a close match to that of the full-order system.

**Figure 15:**
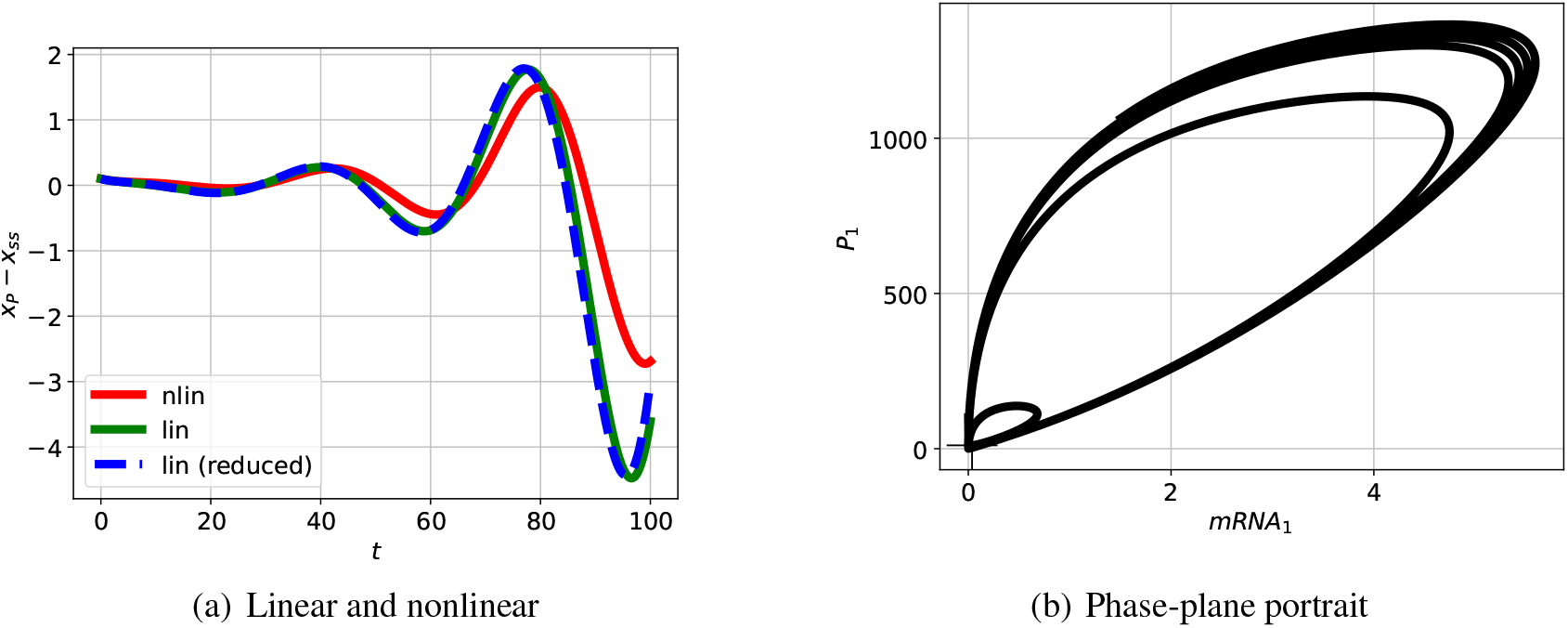
Repressilator: nonlinear simulation. (a) The incremental response of the indicated state to an initial condition corresponding to the perturbed steady-state and plotted against time *t* for a short time span. As expected, the initial nonlinear and linear responses are close for the small initial perturbation; as indicated in (b) the reponses diverge as time increases. (b) The phase-plane response corresponding to two particular states plotted for a longer time. The steady-state value is marked by +. The trajectory spirals outwards from the initial condition to the limit cycle. The simulation terminates at *t* = 250min.

Figure 15(b) shows the nonlinear response when the steady-state is perturbed plotted as a phase-plane diagram. As expected, the initial response is a spiral corresponding to the linear oscillation, but the the response settles to a limit cycle.

For this example, a system with 46 states can be effectively analysed using a reduced-order loop gain transfer function *L* of fourth order.

## 6 Conclusion

Energy-based modelling of biochemical systems using the bond graph approach was combined with linearisation and classical feedback control theory to give a novel approach to the analysis of biochemical oscillators. In particular, a *frequency-domain* approach enabled classical control theory concepts, in particular the Bode diagram, to be used. This allows control engineering insights [47] to be reused in this context.

It was demonstrated that the stability of the linearised system is dependent on the interplay between *active* and *passive* feedback and this interplay is formalised using classical frequency-response analysis of feedback systems. In particular, the *phase margin* was suggested as a simple scalar indicator of the stability of the linearised system. The properties of biochemical systems are dependent on both the structure of the systems as well and the parameters of the individual components. It was shown that investigating the effect of structural and parameter variation on the phase margin provides a convenient and simulation-free approach to the analysis of how system structure and parameters gives rise to oscillations. The resultant design can then be analysed using non-linear techniques [35, 36].

The method was illustrated using three specific examples. However, it is believed that the approach is applicable to a wide range of biological oscillators; this is a topic of current research.

Computing the frequency response of a high order system is straightforward using well-established methods [62] and thus the application of the phase margin approach is scaleable to high-order systems. The repressilator example of § 5 provides an illustration of a moderately high-order system of 46 states. It was shown that further insight into oscillator properties can be found using the root-locus approach and this was illustrated using the examples of §§ 3 & 4. Although convenient for low-order systems, the root-locus approach does not scale to high order systems. However in the case of the repressilator example of § 5, it was found that the relevant properties of the system where expressed by a low order approximation and thus the root-locus method gave further insight in this case. Once again, this reduction was achieved by well-established scaleable methods [62].

As mentioned in the Introduction, a number of previous approaches [5–9] have used linearisation to predict oscillatory behaviour, this paper expands the approach in a number of ways including the use of frequency domain analysis via the Bode diagram and the associated phase margin concept. The division of the feedback loop into active and passive components gives novel insights into the conditions for oscillation and the significance of parametric and structural variation.

Although the methods of this paper are aimed at the *analysis* of biochemical oscillators, it is believed that the insights obtained by the phase margin approach and the concepts of active and passive feedback provide a basis for the *synthesis* of biochemical oscillators. In particular, the active and passive feedback components may point towards the structural and parameter changes required to drive biological systems into oscillatory regimes, building on previous work on tunable synthetic oscillators [64, 65].

The bond graph approach is an energy based method which has previously been used to study the energy consumption of action potentials and synthetic oscillators [15, 16]. The bond graph approach can therefore be used assess the energy requirements of biochemical oscillators as part of the synthesis process; this is a topic of future research. In synthetic biology, excessive energy consumption is a common cause of the failure of novel circuits [66]; we speculate that a bond graph approach can highlight important trade-offs between resource consumption, robustness and performance.

## Supporting information

Reactions, state equations and parameters.

## 7 Author Contributions

PJG prepared the numerical results and drafted the manuscript; PJG and MP discussed the results and revised the manuscript. PJG and MP approved the final version for publication.

## 8 Acknowledgements

PJG would like to thank the Faculty of Engineering and Information Technology, University of Melbourne, for its support via a Professorial Fellowship. The authors would like to thank the anonymous reviewers for their constructive comments which have enhanced the paper.

## Notes

https://github.com/gawthrop/Oscillation24

### Competing Interest Statement

The authors have declared no competing interest.

### Summary of Updates

This paper has been revised to improve clarity. Additional supplementary material has been prepared.

## References

[1] Daniel B Forger. Biological clocks, rhythms, and oscillations: the theory of biological timekeeping. MIT Press, Cambridge Mass., 2017. ISBN 9780262036771.

[2] Brian C. Goodwin. Temporal Organization in Cells. Academic Press, London and New York, 1963.

[3] A. Johnsson and H.G. Karlsson. A feedback model for biological rhythms: I. mathematical description and basic properties of the model. Journal of Theoretical Biology, 36(1):153–174, 1972. doi:10.1016/0022-5193(72)90185-3.

[4] John J. Tyson. Periodic enzyme synthesis: Reconsideration of the theory of oscillatory repression. Journal of Theoretical Biology, 80(1):27–38, 1979. doi:10.1016/0022-5193(79)90177-2.

[5] John J. Tyson. Biochemical oscillations. In Christopher P. Fall, Eric S. Marland, John M. Wagner, and John J. Tyson, editors, Computational Cell Biology, chapter 9, pages 230–260. Springer New York, New York, NY, 2002. ISBN 978-0-387-22459-6. doi:10.1007/978-0-38722459-69.

[6] Luonan Chen and K. Aihara. A model of periodic oscillation for genetic regulatory systems. IEEE Transactions on Circuits and Systems I: Fundamental Theory and Applications, 49(10): 1429–1436, 2002. doi:10.1109/TCSI.2002.803354.

[7] Béla Novák and John J. Tyson. Design principles of biochemical oscillators. Nature Reviews Molecular Cell Biology, 9(12):981–991, Dec 2008. doi:10.1038/nrm2530.

[8] Neslihan Avcu and Cüneyt Güzeliş. Bifurcation analysis of bistable and oscillatory dynamics in biological networks using the root-locus method. IET Systems Biology, 13(6):333–345, 2019. doi:10.1049/iet-syb.2019.0043.

[9] John J. Tyson and Béla Novák. Time-keeping and decision-making in the cell cycle. Interface Focus, 12(4):20210075, 2022. doi:10.1098/rsfs.2021.0075.

[10] Didier Gonze and Peter Ruoff. The Goodwin oscillator and its legacy. Acta Biotheoretica, 69 (4):857–874, Dec 2021. doi:10.1007/s10441-020-09379-8.

[11] E.E. Sel’kov. Self-oscillations in glycolysis 1. a simple kinetic model. European Journal of Biochemistry, 4(1):79 – 86, 1968. doi:10.1111/j.1432-1033.1968.tb00175.x.

[12] James P Keener and James Sneyd. Mathematical Physiology: I: Cellular Physiology, volume 1. Springer, New York, 2nd edition, 2009.

[13] Michael B. Elowitz and Stanislas Leibler. A synthetic oscillatory network of transcriptional regulators. Nature, 403(6767):335–338, Jan 2000. doi:10.1038/35002125.

[14] John J Tyson, Katherine C Chen, and Bela Novak. Sniffers, buzzers, toggles and blinkers: dynamics of regulatory and signaling pathways in the cell. Current Opinion in Cell Biology, 15 (2):221 – 231, 2003. doi:10.1016/S0955-0674(03)00017-6.

[15] Michael Pan, Peter J. Gawthrop, Kenneth Tran, Joseph Cursons, and Edmund J. Crampin. Bond graph modelling of the cardiac action potential: implications for drift and non-unique steady states. Proceedings of the Royal Society A: Mathematical, Physical and Engineering Sciences, 474(2214):20180106, Jun 2018. doi:10.1098/rspa.2018.0106.

[16] Michael Pan, Peter J. Gawthrop, Matthew Faria, and Stuart T. Johnston. Thermodynamically-consistent, reduced models of gene regulatory networks. bioRxiv, 2023. doi:10.1101/2023.11.13.566770.

[17] H. M. Paynter. Analysis and Design of Engineering Systems. MIT Press, Cambridge, Mass., 1961.

[18] P. J. Gawthrop and L. P. S. Smith. Metamodelling: Bond Graphs and Dynamic Systems. Prentice Hall, Hemel Hempstead, Herts, England., 1996. ISBN 0-13-489824-9. doi:10.5281/zenodo.6998395.

[19] Peter J Gawthrop and Geraint P Bevan. Bond-graph modeling: A tutorial introduction for control engineers. IEEE Control Systems Magazine, 27(2):24–45, April 2007. doi:10.1109/MCS.2007.338279.

[20] Dean C Karnopp, Donald L Margolis, and Ronald C Rosenberg. System Dynamics: Modeling, Simulation, and Control of Mechatronic Systems. John Wiley & Sons, Hoboken, New Jersey, 5th edition, 2012. ISBN 978-0470889084.

[21] Wolfgang Borutzky, editor. Bond Graphs for Modelling, Control and Fault Diagnosis of Engineering Systems. Springer New York, 2017. doi:10.1007/978-3-319-47434-2. In press.

[22] George Oster, Alan Perelson, and Aharon Katchalsky. Network thermodynamics. Nature, 234: 393–399, December 1971. doi:10.1038/234393a0.

[23] George F. Oster, Alan S. Perelson, and Aharon Katchalsky. Network thermodynamics: dynamic modelling of biophysical systems. Quarterly Reviews of Biophysics, 6(01):1–134, 1973. doi:10.1017/S0033583500000081.

[24] Peter J. Gawthrop and Edmund J. Crampin. Energy-based analysis of biochemical cycles using bond graphs. Proceedings of the Royal Society A: Mathematical, Physical and Engineering Science, 470(2171):1–25, 2014. doi:10.1098/rspa.2014.0459. Available at 1406.2447.

[25] Peter J. Gawthrop and Michael Pan. Network thermodynamics of biological systems: A bond graph approach. Mathematical Biosciences, 352:108899, 2022. doi:10.1016/j.mbs.2022.108899.

[26] Peter Hunter. The virtual physiological human: The physiome project aims to develop reproducible, multiscale models for clinical practice. IEEE Pulse, 7(4):36–42, July 2016. ISSN 2154-2287. doi:10.1109/MPUL.2016.2563841.

[27] Vijay Rajagopal, Senthil Arumugam, Peter J. Hunter, Afshin Khadangi, Joshua Chung, and Michael Pan. The cell physiome: What do we need in a computational physiology framework for predicting single-cell biology? Annual Review of Biomedical Data Science, 5(1):341–366, 2022. doi:10.1146/annurev-biodatasci-072018-021246. PMID: 35576556.

[28] P. J. Gawthrop and E. J. Crampin. Modular bond-graph modelling and analysis of biomolecular systems. IET Systems Biology, 10(5):187–201, October 2016. doi:10.1049/iet-syb.2015.0083. Available at 1511.06482.

[29] Peter J. Gawthrop, Joseph Cursons, and Edmund J. Crampin. Hierarchical bond graph modelling of biochemical networks. Proceedings of the Royal Society A: Mathematical, Physical and Engineering Sciences, 471(2184):1–23, 2015. doi:10.1098/rspa.2015.0642.

[30] Michael Pan, Peter J. Gawthrop, Joseph Cursons, and Edmund J. Crampin. Modular assembly of dynamic models in systems biology. PLOS Computational Biology, 17(10):e1009513, Oct 2021. doi:10.1371/journal.pcbi.1009513.

[31] Peter J. Gawthrop, Michael Pan, and Edmund J. Crampin. Modular dynamic biomolecular modelling with bond graphs: the unification of stoichiometry, thermodynamics, kinetics and data. Journal of The Royal Society Interface, 18(181):20210478, 2021. doi:10.1098/rsif.2021.0478.

[32] Stefano Stramigioli and Michel van Dijk. Energy conservative limit cycle oscillations. IFAC Proceedings Volumes, 41(2):15666 – 15671, 2008. doi:10.3182/20080706-5-KR-1001.02649. 17th IFAC World Congress.

[33] Martin Feinberg. Foundations of Chemical Reaction Network Theory. Springer, 2019. ISBN 978-3-030-03857-1. doi:10.1007/978-3-030-03858-8.

[34] P. Gawthrop and E. J. Crampin. Bond graph representation of chemical reaction networks. IEEE Transactions on NanoBioscience, 17(4):449–455, October 2018. ISSN 1536-1241. doi:10.1109/TNB.2018.2876391. Available at 1809.00449.

[35] Dominic Jordan and Peter Smith. Nonlinear Ordinary Differential Equations: An Introduction for Scientists and Engineers. Oxford University Press, Oxford, 2007.

[36] M.W. Hirsch, S. Smale, and R.L. Devaney. Differential Equations, Dynamical Systems, and an Introduction to Chaos. Academic Press, third edition, 2012. ISBN 978-0-12-382010-5.

[37] Charles F. Walter. The occurrence and the significance of limit cycle behavior in controlled biochemical systems. Journal of Theoretical Biology, 27(2):259–272, 1970. doi:10.1016/0022-5193(70)90141-4.

[38] Albert Goldbeter. Biochemical Oscillations and Cellular Rhythms: the Molecular Bases of Periodic and Chaotic Behaviour. Cambridge University Press, 1996. ISBN 0 521 40307 3.

[39] Claude Gérard and Albert Goldbeter. The cell cycle is a limit cycle. Mathematical Modelling of Natural Phenomena, 7(6):126–166, 2012.

[40] Aurore Woller, Didier Gonze, and Thomas Erneux. The Goodwin model revisited: Hopf bifurcation, limit-cycle, and periodic entrainment. Physical Biology, 11(4):045002, Jul 2014. doi:10.1088/1478-3975/11/4/045002.

[41] Vitaly A. Likhoshvai, Vladimir P. Golubyatnikov, and Tamara M. Khlebodarova. Limit cycles in models of circular gene networks regulated by negative feedback loops. BMC Bioinformatics, 21(11):255, Sep 2020. doi:10.1186/s12859-020-03598-z.

[42] Sandip Saha, Gautam Gangopadhyay, and Deb Shankar Ray. Universality in bio-rhythms: A perspective from nonlinear dynamics. Journal of Biosciences, 47(1):16, Mar 2022. doi:10.1007/s12038-021-00249-0.

[43] Arthur Gelb and Wallace E. Vander Velde. Multiple-input describing functions and nonlinear system design. McGraw Hill, New York, 1968.

[44] O. L. R. Jacobs. Introduction to Control Theory. Oxford University Press, 1974.

[45] D.P. Atherton. Early developments in nonlinear control. IEEE Control Systems Magazine, 16 (3):34–43, June 1996. doi:10.1109/37.506396.

[46] Derek P Atherton. An introduction to nonlinearity in control systems. Bookboon, 2011.

[47] Karl Johan Åström and Richard M Murray. Feedback systems: an introduction for scientists and engineers. Princeton University Press, 2008. ISBN 978-0-691-13576-2.

[48] Ion Victor Gosea. Exact and inexact lifting transformations of nonlinear dynamical systems: Transfer functions, equivalence, and complexity reduction. Applied Sciences, 12(5), 2022. ISSN 2076-3417. doi:10.3390/app12052333.

[49] P. J. Gawthrop. Energy-based modeling of the feedback control of biomolecular systems with cyclic flow modulation. IEEE Transactions on NanoBioscience, 20(2):183–192, April 2021. doi:10.1109/TNB.2021.3058440.

[50] Dean Karnopp. Bond graphs in control: Physical state variables and observers. Journal of the Franklin Institute, 308(3):219 – 234, 1979. doi:10.1016/0016-0032(79)90114-5.

[51] N. Hogan. Impedance control: An approach to manipulation. part I–theory. ASME Journal of Dynamic Systems, Measurement and Control, 107:1–7, March 1985.

[52] A. Sharon, N. Hogan, and D. E. Hardt. Controller design in the physical domain. Journal of the Franklin Institute, 328(5):697–721, 1991.

[53] Peter J. Gawthrop. Physical model-based control: A bond graph approach. Journal of the Franklin Institute, 332(3):285–305, 1995. doi:10.1016/0016-0032(95)00044-5.

[54] D.J. Costello and P.J. Gawthrop. Physical-model based control: Experiments with a stirred-tank heater. Chemical Engineering Research and Design, 75(3):361–370, 1997. doi:10.1205/026387697523679. Particle Processing.

[55] P.J. Gawthrop, B. Bhikkaji, and S.O.R. Moheimani. Physical-model-based control of a piezoelectric tube for nano-scale positioning applications. Mechatronics, 20(1):74 – 84, February 2010. doi:10.1016/j.mechatronics.2009.09.006. Available online 13 October 2009.

[56] Peter Gawthrop, S.A. Neild, and D.J. Wagg. Dynamically dual vibration absorbers: a bond graph approach to vibration control. Systems Science and Control Engineering, 3(1):113–128, 2015. doi:10.1080/21642583.2014.991458.

[57] Domitilla Del Vecchio, Alexander J. Ninfa, and Eduardo D. Sontag. Modular cell biology: retroactivity and insulation. Molecular Systems Biology, 4:1–16, 2008. doi:10.1038/msb4100204.

[58] Domitilla Del Vecchio and Eduardo D. Sontag. Engineering principles in bio-molecular systems: From retroactivity to modularity. European Journal of Control, 15(3–4):389 – 397, 2009. doi:10.3166/ejc.15.389-397.

[59] Domitilla Del Vecchio and Richard M Murray. Biomolecular Feedback Systems. Princeton University Press, 2014. ISBN 0691161534.

[60] Jeffrey M Perkel. By Jupyter, it all makes sense. Nature, 563(7729):145–146, 2018.

[61] Peter Cudmore, Michael Pan, Peter J. Gawthrop, and Edmund J. Crampin. Analysing and simulating energy-based models in biology using BondGraphTools. The European Physical Journal E, 44(12):148, Dec 2021. doi:10.1140/epje/s10189-021-00152-4.

[62] Sawyer Fuller, Ben Greiner, Jason Moore, Richard Murray, René van Paassen, and Rory Yorke. The python control systems library (python-control). In 2021 60th IEEE Conference on Decision and Control (CDC), pages 4875–4881, Austin, Texas, Dec 2021. doi:10.1109/CDC45484.2021.9683368.

[63] John J Tyson, Reka Albert, Albert Goldbeter, Peter Ruoff, and Jill Sible. Biological switches and clocks. Journal of The Royal Society Interface, 5:S1–S8, 2008. doi:10.1098/rsif.2008.0179.focus.

[64] Jesse Stricker, Scott Cookson, Matthew R. Bennett, William H. Mather, Lev S. Tsimring, and Jeff Hasty. A fast, robust and tunable synthetic gene oscillator. Nature, 456(7221):516–519, Nov 2008. ISSN 1476-4687. doi:10.1038/nature07389.

[65] Marcel Tigges, Tatiana T. Marquez-Lago, Jörg Stelling, and Martin Fussenegger. A tunable synthetic mammalian oscillator. Nature, 457(7227):309–312, Jan 2009. ISSN 1476-4687. doi:10.1038/nature07616.

[66] Andrea Y. Weiße, Diego A. Oyarzún, Vincent Danos, and Peter S. Swain. Mechanistic links between cellular trade-offs, gene expression, and growth. Proceedings of the National Academy of Sciences, 112(9):E1038–E1047, 2015. ISSN 0027-8424. doi:10.1073/pnas.1416533112.

