## Additional figures. for "Energy-based Analysis of Biochemical Oscillators Using Bond Graphs and Linear Control Theory"

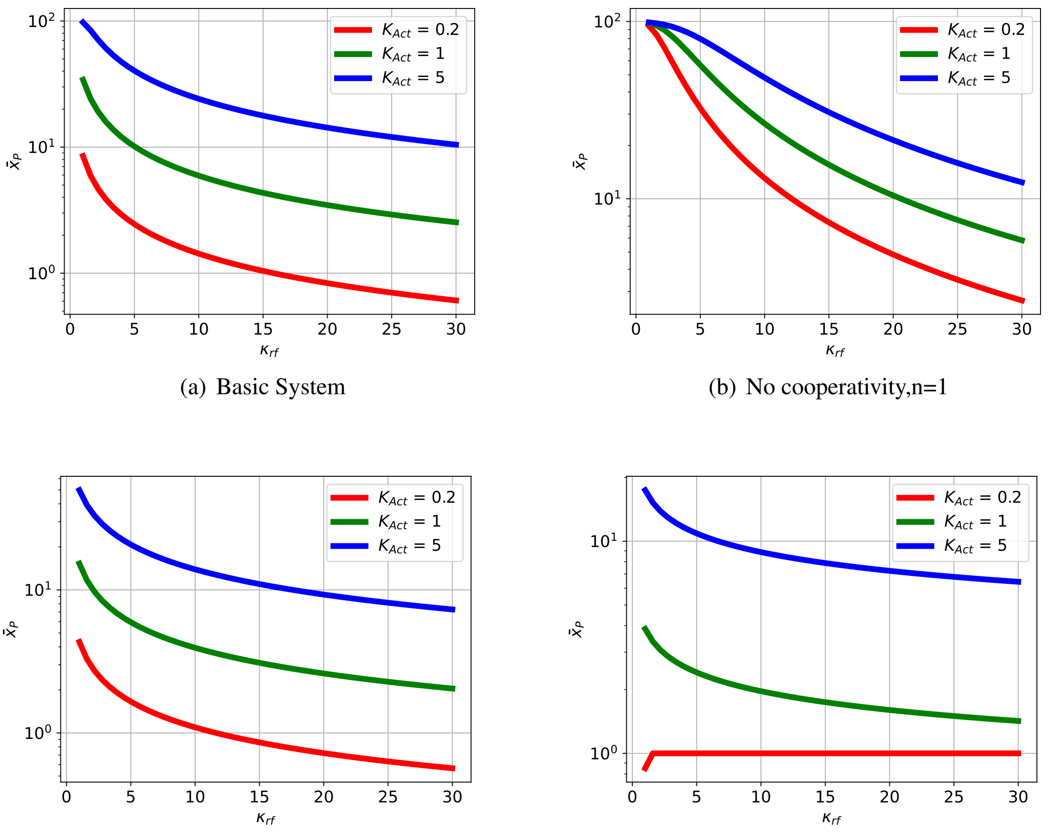


(a) Basic System

(b) No cooperativity,n=1

(c) N=2

(d) N=2, enhanced cooperativity, n=4

Figure S1: Illustrative Example.The steady-state value $\bar{x}_P$ of $x_P$ as parameters vary for three systems with modified structure.The system steady state $\bar{x}_P$ (about which the system is linearised) is dependent on both system parameters and structure.

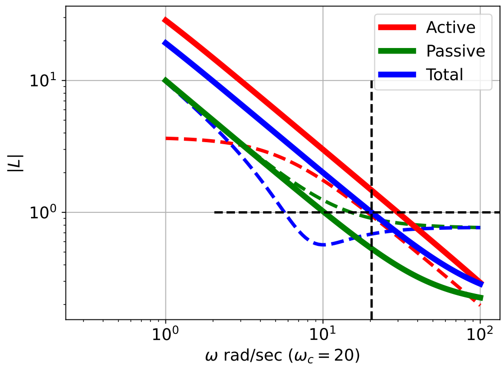


(a) Bode diagram: gain

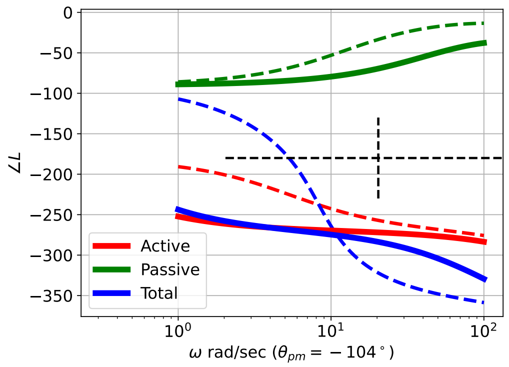


(b) Bode diagram: phase

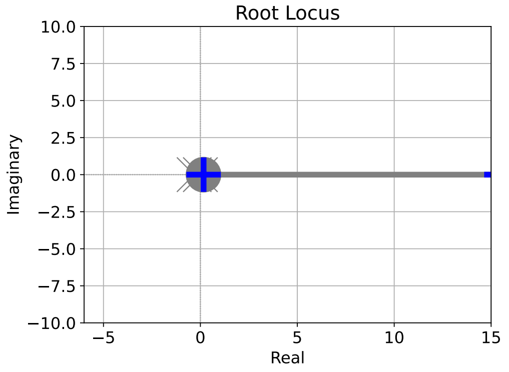


(c) Root-locus diagram

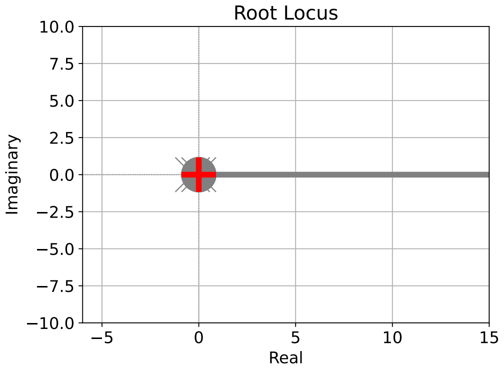


(d) Root-locus diagram (active)

Figure S2: Sel'kov Glycolytic Oscillator: extra cooperativity: comparative analysis. (a)&(b) As 3(a) & 3(b). In this case, the phase margin is $\theta_{pm} = -104^\circ$ at $\omega_{pm} = 20\,\mathrm{rad\,s^{-1}}$ ; the negative sign indicating instability. In this case, the passive part $L_{Pas}(s)$ is negligible and the closed loop system behaves as a typical positive-feedback system corresponding to the active part $L_{Act}(s)$ with an increasing exponential response. The closed-loop poles a real indicating no oscillatory component. (c)&(d) As 3(c) & 3(d).

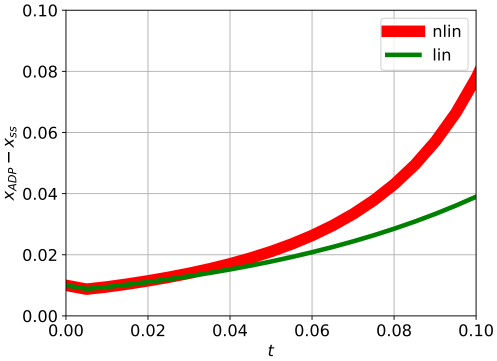


(a) Linear and nonlinear

Figure S3: Sel'kov Glycolytic Oscillator: extra cooperativity: nonlinear simulation. The trajectory is an increasing exponential.
