## Supplementary material for "Energy-based Analysis of Biochemical Oscillators Using Bond Graphs and Linear Control Theory": Reactions, state equations and parameters.

October 11, 2024

### 1 Introduction

This Jupyter notebook (Supplementary.ipynb) prints state equations and parameters for the three examples of the paper “Analysis of Biochemical Oscillators Using Bond Graphs and Linear Control Theory” by Peter Gawthrop and Michael Pan:

- 3. Illustrative Example (system Toy)
- 4. The Sel’kov Oscillator (system Selkov)
- 5. The Repressilator

The state equations are automatically generated from the bond graph. Following standard systems biology practice, the state equations are shown in two parts:

- The state derivatives $\dot{x}$ in terms of the reaction flows $v$.
- The reaction flows $v$ in terms of the states $x$.

[1]:
```
# Display LaTeX
import IPython.display as disp

## Stoichiometric analysis
import stoich as st

## Data files
import pickle

# Allow output from within functions
from IPython.core.interactiveshell import InteractiveShell
InteractiveShell.ast_node_interactivity = "all"

chemformula = False
split = 10
```

### 2 System Toy

[2]:
```
## Get data
SysName = 'Toy'
file = open(f'{SysName}.dat', 'rb')
SavedData = pickle.load(file)
file.close()

Stoich = SavedData['Stoich']
s = Stoich['s']
sc = Stoich['sc']
parameter = Stoich['parameter']
```

#### 2.1 List of Reactions

[3]:
```
## Reactions
disp.Latex(st.sprintrl(s,chemformula=chemformula,split=10,all=True))
```
[3]:

$$fbAct + fbE0 \Leftrightarrow E1 + P \tag{2.1.1}$$
$$2E1 + decr1A \Leftrightarrow 2E1 + E2 \tag{2.1.2}$$
$$E2 \Leftrightarrow decr1Zf \tag{2.1.3}$$
$$2E2 + decr2A \Leftrightarrow 2E2 + E3 \tag{2.1.4}$$
$$E3 \Leftrightarrow decr2Zf \tag{2.1.5}$$
$$2E3 + decr3A \Leftrightarrow 2E3 + P \tag{2.1.6}$$
$$P \Leftrightarrow decr3Zf \tag{2.1.7}$$

#### 2.2 List of Flows $v$ as Function of States $x$

[4]:
```
## Flows
disp.Latex(st.sprintvl(s))
```
[4]:

$$v_{fbr} = \kappa_{fbr}\left(-K_{E1}K_{P}x_{E1}x_{P} + K_{fbAct}K_{fbE0}x_{fbAct}x_{fbE0}\right) \tag{2.2.1}$$
$$v_{decr1r} = K_{E1}^{2}\kappa_{decr1r}x_{E1}^{2}\left(-K_{E2}x_{E2} + K_{decr1A}x_{decr1A}\right) \tag{2.2.2}$$
$$v_{decr1rf} = \kappa_{decr1rf}\left(K_{E2}x_{E2} - K_{decr1Zf}x_{decr1Zf}\right) \tag{2.2.3}$$
$$v_{decr2r} = K_{E2}^{2}\kappa_{decr2r}x_{E2}^{2}\left(-K_{E3}x_{E3} + K_{decr2A}x_{decr2A}\right) \tag{2.2.4}$$
$$v_{decr2rf} = \kappa_{decr2rf}\left(K_{E3}x_{E3} - K_{decr2Zf}x_{decr2Zf}\right) \tag{2.2.5}$$
$$v_{decr3r} = K_{E3}^{2}\kappa_{decr3r}x_{E3}^{2}\left(-K_{P}x_{P} + K_{decr3A}x_{decr3A}\right) \tag{2.2.6}$$
$$v_{decr3rf} = \kappa_{decr3rf}\left(K_{P}x_{P} - K_{decr3Zf}x_{decr3Zf}\right) \tag{2.2.7}$$

#### 2.3 List of State Derivatives $\dot{x}$ as Function of Flows $v$.

[5]:
```
## State equations
eqns = st.sprintdxl(s,sc)
disp.Latex(eqns)
```
[5]:

$$\dot{x}_{E1} = v_{fbr} \tag{2.3.1}$$
$$\dot{x}_{E2} = v_{decr1r} - v_{decr1rf} \tag{2.3.2}$$
$$\dot{x}_{E3} = v_{decr2r} - v_{decr2rf} \tag{2.3.3}$$

#### 2.4 List of Parameters.

[6]:
```
## Parameters
pars = st.sprintparl(parameter)
disp.Latex(pars)
```

[6]:

$$K_{fbAct} = 1 \quad (2.4.1)$$
$$K_{decr1Zf} = 1e-06 \quad (2.4.2)$$
$$\kappa_{decr1r} = 1 \quad (2.4.3)$$
$$\kappa_{decr1rf} = 10 \quad (2.4.4)$$
$$K_{decr1A} = 100 \quad (2.4.5)$$
$$K_{E1} = 1 \quad (2.4.6)$$
$$K_{decr2Zf} = 1e-06 \quad (2.4.7)$$
$$\kappa_{decr2r} = 1 \quad (2.4.8)$$
$$\kappa_{decr2rf} = 10 \quad (2.4.9)$$
$$K_{decr2A} = 100 \quad (2.4.10)$$

$$K_{E2} = 1 \quad (2.4.11)$$
$$K_{decr3Zf} = 1e-06 \quad (2.4.12)$$
$$\kappa_{decr3r} = 1 \quad (2.4.13)$$
$$\kappa_{decr3rf} = 10 \quad (2.4.14)$$
$$K_{decr3A} = 100 \quad (2.4.15)$$
$$K_{E3} = 1 \quad (2.4.16)$$

### 3 System Selkov

[7]:
```
## Get data
SysName = 'Selkov'
file = open(f'{SysName}.dat', 'rb')
SavedData = pickle.load(file)
file.close()

Stoich = SavedData['Stoich']
s = Stoich['s']
sc = Stoich['sc']
parameter = Stoich['parameter']
```

#### 3.1 List of Reactions

[8]:
```
## Reactions
disp.Latex(st.sprintrl(s,chemformula=chemformula,all=True))
```

[8]:

$$2P + selkovPFK \Leftrightarrow selkovE \tag{3.1.1}$$
$$selkovATP + selkovE \Leftrightarrow selkovC \tag{3.1.2}$$
$$selkovC \Leftrightarrow P + selkovE + selkovZ \tag{3.1.3}$$
$$P \Leftrightarrow selkovZf \tag{3.1.4}$$
$$selkovATP0 \Leftrightarrow selkovATP \tag{3.1.5}$$

#### 3.2 List of Flows $v$ as Function of States $x$

[9]:
```
## Flows
disp.Latex(st.sprintvl(s))
```

[9]:

$$v_{selkovr0} = \kappa_{selkovr0} \left(K_P^2 K_{selkovPFK} x_P^2 x_{selkovPFK} - K_{selkovE} x_{selkovE}\right) \tag{3.2.1}$$
$$v_{selkovr1} = \kappa_{selkovr1} \left(K_{selkovATP} K_{selkovE} x_{selkovATP} x_{selkovE} - K_{selkovC} x_{selkovC}\right) \tag{3.2.2}$$
$$v_{selkovr2} = \kappa_{selkovr2} \left(-K_P K_{selkovE} K_{selkovZ} x_P x_{selkovE} x_{selkovZ} + K_{selkovC} x_{selkovC}\right) \tag{3.2.3}$$
$$v_{selkovrf} = \kappa_{selkovrf} \left(K_P x_P - K_{selkovZf} x_{selkovZf}\right) \tag{3.2.4}$$
$$v_{selkovrs} = \kappa_{selkovrs} \left(-K_{selkovATP} x_{selkovATP} + K_{selkovATP0} x_{selkovATP0}\right) \tag{3.2.5}$$

#### 3.3 List of State Derivatives $\dot{x}$ as Function of Flows $v$.

[10]:
```
## State equations
eqns = st.sprintdxl(s,sc)
disp.Latex(eqns)
```

[10]:

$$\dot{x}_{selkovATP} = -v_{selkovr1} + v_{selkovrs} \tag{3.3.1}$$
$$\dot{x}_{selkovC} = v_{selkovr1} - v_{selkovr2} \tag{3.3.2}$$
$$\dot{x}_{selkovE} = v_{selkovr0} - v_{selkovr1} + v_{selkovr2} \tag{3.3.3}$$
$$\dot{x}_{selkovPFK} = -v_{selkovr0} \tag{3.3.4}$$

#### 3.4 List of Parameters.

[11]:
```
## Parameters
pars = st.sprintparl(parameter)
disp.Latex(pars)
```

[11]:

$$K_{selkovZ} = 1e-10 \tag{3.4.1}$$
$$K_{selkovZf} = 1e-10 \tag{3.4.2}$$
$$\kappa_{selkovr0} = 1000 \tag{3.4.3}$$
$$\kappa_{selkovr1} = 1000 \tag{3.4.4}$$
$$\kappa_{selkovr2} = 1000 \tag{3.4.5}$$
$$\kappa_{selkovrf} = 10 \tag{3.4.6}$$
$$K_{selkovPFK} = 1 \tag{3.4.7}$$
$$K_{selkovC} = 1 \tag{3.4.8}$$
$$K_{selkovATP0} = 1000 \tag{3.4.9}$$
$$\kappa_{selkovrs} = 0.0006 \tag{3.4.10}$$

### 4 System Repressilator

[12]:
```
## Get data
SysName = 'Repressilator'
file = open(f'{SysName}.dat', 'rb')
SavedData = pickle.load(file)
file.close()

Stoich = SavedData['Stoich']
s = Stoich['s']
sc = Stoich['sc']
parameter = Stoich['parameter']
```

#### 4.1 List of Reactions

[13]:
```
## Reactions
disp.Latex(st.sprintrl(s,chemformula=chemformula,split=split,all=True))
```

[13]:

$$A + G1E \Leftrightarrow G1EA \tag{4.1.1}$$
$$G1EA \Leftrightarrow G1E + G1M \tag{4.1.2}$$
$$G1E + 2P2 \Leftrightarrow G1EI \tag{4.1.3}$$
$$G1EA + 2P2 \Leftrightarrow G1EAI \tag{4.1.4}$$
$$G1M + R \Leftrightarrow G1C0 \tag{4.1.5}$$
$$A + G1C0 \Leftrightarrow G1M + G1C1 \tag{4.1.6}$$
$$A + G1C1 \Leftrightarrow G1C2 \tag{4.1.7}$$
$$A + G1C2 \Leftrightarrow G1C3 \tag{4.1.8}$$
$$A + G1C3 \Leftrightarrow G1C4 \tag{4.1.9}$$
$$A + G1C4 \Leftrightarrow G1C5 \tag{4.1.10}$$

$$A + G1C5 \Leftrightarrow G1C6 \tag{4.1.11}$$
$$A + G1C6 \Leftrightarrow G1C7 \tag{4.1.12}$$
$$A + G1C7 \Leftrightarrow G1C8 \tag{4.1.13}$$
$$G1C8 \Leftrightarrow R + P1 \tag{4.1.14}$$
$$G1M \Leftrightarrow G1XM \tag{4.1.15}$$
$$P1 \Leftrightarrow G1XP \tag{4.1.16}$$
$$A + G2E \Leftrightarrow G2EA \tag{4.1.17}$$
$$G2EA \Leftrightarrow G2E + G2M \tag{4.1.18}$$
$$G2E + 2P3 \Leftrightarrow G2EI \tag{4.1.19}$$
$$G2EA + 2P3 \Leftrightarrow G2EAI \tag{4.1.20}$$

$$R + G2M \Leftrightarrow G2C0 \tag{4.1.21}$$
$$A + G2C0 \Leftrightarrow G2M + G2C1 \tag{4.1.22}$$
$$A + G2C1 \Leftrightarrow G2C2 \tag{4.1.23}$$
$$A + G2C2 \Leftrightarrow G2C3 \tag{4.1.24}$$
$$A + G2C3 \Leftrightarrow G2C4 \tag{4.1.25}$$
$$A + G2C4 \Leftrightarrow G2C5 \tag{4.1.26}$$
$$A + G2C5 \Leftrightarrow G2C6 \tag{4.1.27}$$
$$A + G2C6 \Leftrightarrow G2C7 \tag{4.1.28}$$
$$A + G2C7 \Leftrightarrow G2C8 \tag{4.1.29}$$
$$G2C8 \Leftrightarrow P2 + R \tag{4.1.30}$$

$$G2M \Leftrightarrow G2XM \tag{4.1.31}$$
$$P2 \Leftrightarrow G2XP \tag{4.1.32}$$
$$A + G3E \Leftrightarrow G3EA \tag{4.1.33}$$
$$G3EA \Leftrightarrow G3E + G3M \tag{4.1.34}$$
$$2P1 + G3E \Leftrightarrow G3EI \tag{4.1.35}$$
$$2P1 + G3EA \Leftrightarrow G3EAI \tag{4.1.36}$$
$$R + G3M \Leftrightarrow G3C0 \tag{4.1.37}$$
$$A + G3C0 \Leftrightarrow G3M + G3C1 \tag{4.1.38}$$
$$A + G3C1 \Leftrightarrow G3C2 \tag{4.1.39}$$
$$A + G3C2 \Leftrightarrow G3C3 \tag{4.1.40}$$

$$A + G3C3 \Leftrightarrow G3C4 \tag{4.1.41}$$
$$A + G3C4 \Leftrightarrow G3C5 \tag{4.1.42}$$
$$A + G3C5 \Leftrightarrow G3C6 \tag{4.1.43}$$
$$A + G3C6 \Leftrightarrow G3C7 \tag{4.1.44}$$
$$A + G3C7 \Leftrightarrow G3C8 \tag{4.1.45}$$
$$G3C8 \Leftrightarrow R + P3 \tag{4.1.46}$$
$$G3M \Leftrightarrow G3XM \tag{4.1.47}$$
$$P3 \Leftrightarrow G3XP \tag{4.1.48}$$

#### 4.2 List of Flows $v$ as Function of States $x$

[14]:
```
## Flow
disp.Latex(st.sprintvl(s,split=10))
```

[14]:

$$v_{G1Tc1} = \kappa_{G1Tc1} \left(K_A K_{G1E} x_A x_{G1E} - K_{G1EA} x_{G1EA}\right) \tag{4.2.1}$$
$$v_{G1Tc2} = \kappa_{G1Tc2} \left(-K_{G1E} K_{G1M} x_{G1E} x_{G1M} + K_{G1EA} x_{G1EA}\right) \tag{4.2.2}$$
$$v_{G1Tc3} = \kappa_{G1Tc3} \left(K_{G1E} K_{P2}^2 x_{G1E} x_{P2}^2 - K_{G1EI} x_{G1EI}\right) \tag{4.2.3}$$
$$v_{G1Tc4} = \kappa_{G1Tc4} \left(K_{G1EA} K_{P2}^2 x_{G1EA} x_{P2}^2 - K_{G1EAI} x_{G1EAI}\right) \tag{4.2.4}$$
$$v_{G1rb} = \kappa_{G1rb} \left(-K_{G1C0} x_{G1C0} + K_{G1M} K_R x_{G1M} x_R\right) \tag{4.2.5}$$
$$v_{G1r1} = \kappa_{G1r1} \left(K_A K_{G1C0} x_A x_{G1C0} - K_{G1C1} K_{G1M} x_{G1C1} x_{G1M}\right) \tag{4.2.6}$$
$$v_{G1r2} = \kappa_{G1r2} \left(K_A K_{G1C1} x_A x_{G1C1} - K_{G1C2} x_{G1C2}\right) \tag{4.2.7}$$
$$v_{G1r3} = \kappa_{G1r3} \left(K_A K_{G1C2} x_A x_{G1C2} - K_{G1C3} x_{G1C3}\right) \tag{4.2.8}$$
$$v_{G1r4} = \kappa_{G1r4} \left(K_A K_{G1C3} x_A x_{G1C3} - K_{G1C4} x_{G1C4}\right) \tag{4.2.9}$$
$$v_{G1r5} = \kappa_{G1r5} \left(K_A K_{G1C4} x_A x_{G1C4} - K_{G1C5} x_{G1C5}\right) \tag{4.2.10}$$

$$v_{G1r6} = \kappa_{G1r6} \left(K_A K_{G1C5} x_A x_{G1C5} - K_{G1C6} x_{G1C6}\right) \tag{4.2.11}$$
$$v_{G1r7} = \kappa_{G1r7} \left(K_A K_{G1C6} x_A x_{G1C6} - K_{G1C7} x_{G1C7}\right) \tag{4.2.12}$$
$$v_{G1r8} = \kappa_{G1r8} \left(K_A K_{G1C7} x_A x_{G1C7} - K_{G1C8} x_{G1C8}\right) \tag{4.2.13}$$
$$v_{G1rt} = \kappa_{G1rt} \left(K_{G1C8} x_{G1C8} - K_{P1} K_R x_{P1} x_R\right) \tag{4.2.14}$$
$$v_{G1degM} = \kappa_{G1degM} \left(K_{G1M} x_{G1M} - K_{G1XM} x_{G1XM}\right) \tag{4.2.15}$$
$$v_{G1degP} = \kappa_{G1degP} \left(-K_{G1XP} x_{G1XP} + K_{P1} x_{P1}\right) \tag{4.2.16}$$
$$v_{G2Tc1} = \kappa_{G2Tc1} \left(K_A K_{G2E} x_A x_{G2E} - K_{G2EA} x_{G2EA}\right) \tag{4.2.17}$$
$$v_{G2Tc2} = \kappa_{G2Tc2} \left(-K_{G2E} K_{G2M} x_{G2E} x_{G2M} + K_{G2EA} x_{G2EA}\right) \tag{4.2.18}$$
$$v_{G2Tc3} = \kappa_{G2Tc3} \left(K_{G2E} K_{P3}^2 x_{G2E} x_{P3}^2 - K_{G2EI} x_{G2EI}\right) \tag{4.2.19}$$
$$v_{G2Tc4} = \kappa_{G2Tc4} \left(K_{G2EA} K_{P3}^2 x_{G2EA} x_{P3}^2 - K_{G2EAI} x_{G2EAI}\right) \tag{4.2.20}$$

$$v_{G2rb} = \kappa_{G2rb} \left(- K_{G2C0} x_{G2C0} + K_{G2M} K_{R} x_{G2M} x_{R}\right) \tag{4.2.21}$$
$$v_{G2r1} = \kappa_{G2r1} \left(K_{A} K_{G2C0} x_{A} x_{G2C0} - K_{G2C1} K_{G2M} x_{G2C1} x_{G2M}\right) \tag{4.2.22}$$
$$v_{G2r2} = \kappa_{G2r2} \left(K_{A} K_{G2C1} x_{A} x_{G2C1} - K_{G2C2} x_{G2C2}\right) \tag{4.2.23}$$
$$v_{G2r3} = \kappa_{G2r3} \left(K_{A} K_{G2C2} x_{A} x_{G2C2} - K_{G2C3} x_{G2C3}\right) \tag{4.2.24}$$
$$v_{G2r4} = \kappa_{G2r4} \left(K_{A} K_{G2C3} x_{A} x_{G2C3} - K_{G2C4} x_{G2C4}\right) \tag{4.2.25}$$
$$v_{G2r5} = \kappa_{G2r5} \left(K_{A} K_{G2C4} x_{A} x_{G2C4} - K_{G2C5} x_{G2C5}\right) \tag{4.2.26}$$
$$v_{G2r6} = \kappa_{G2r6} \left(K_{A} K_{G2C5} x_{A} x_{G2C5} - K_{G2C6} x_{G2C6}\right) \tag{4.2.27}$$
$$v_{G2r7} = \kappa_{G2r7} \left(K_{A} K_{G2C6} x_{A} x_{G2C6} - K_{G2C7} x_{G2C7}\right) \tag{4.2.28}$$
$$v_{G2r8} = \kappa_{G2r8} \left(K_{A} K_{G2C7} x_{A} x_{G2C7} - K_{G2C8} x_{G2C8}\right) \tag{4.2.29}$$
$$v_{G2rt} = \kappa_{G2rt} \left(K_{G2C8} x_{G2C8} - K_{P2} K_{R} x_{P2} x_{R}\right) \tag{4.2.30}$$

$$v_{G2degM} = \kappa_{G2degM} \left(K_{G2M} x_{G2M} - K_{G2XM} x_{G2XM}\right) \tag{4.2.31}$$
$$v_{G2degP} = \kappa_{G2degP} \left(- K_{G2XP} x_{G2XP} + K_{P2} x_{P2}\right) \tag{4.2.32}$$
$$v_{G3Tc1} = \kappa_{G3Tc1} \left(K_{A} K_{G3E} x_{A} x_{G3E} - K_{G3EA} x_{G3EA}\right) \tag{4.2.33}$$
$$v_{G3Tc2} = \kappa_{G3Tc2} \left(- K_{G3E} K_{G3M} x_{G3E} x_{G3M} + K_{G3EA} x_{G3EA}\right) \tag{4.2.34}$$
$$v_{G3Tc3} = \kappa_{G3Tc3} \left(K_{G3E} K_{P1}^{2} x_{G3E} x_{P1}^{2} - K_{G3EI} x_{G3EI}\right) \tag{4.2.35}$$
$$v_{G3Tc4} = \kappa_{G3Tc4} \left(K_{G3EA} K_{P1}^{2} x_{G3EA} x_{P1}^{2} - K_{G3EAI} x_{G3EAI}\right) \tag{4.2.36}$$
$$v_{G3rb} = \kappa_{G3rb} \left(- K_{G3C0} x_{G3C0} + K_{G3M} K_{R} x_{G3M} x_{R}\right) \tag{4.2.37}$$
$$v_{G3r1} = \kappa_{G3r1} \left(K_{A} K_{G3C0} x_{A} x_{G3C0} - K_{G3C1} K_{G3M} x_{G3C1} x_{G3M}\right) \tag{4.2.38}$$
$$v_{G3r2} = \kappa_{G3r2} \left(K_{A} K_{G3C1} x_{A} x_{G3C1} - K_{G3C2} x_{G3C2}\right) \tag{4.2.39}$$
$$v_{G3r3} = \kappa_{G3r3} \left(K_{A} K_{G3C2} x_{A} x_{G3C2} - K_{G3C3} x_{G3C3}\right) \tag{4.2.40}$$

$$v_{G3r4} = \kappa_{G3r4} \left(K_{A} K_{G3C3} x_{A} x_{G3C3} - K_{G3C4} x_{G3C4}\right) \tag{4.2.41}$$
$$v_{G3r5} = \kappa_{G3r5} \left(K_{A} K_{G3C4} x_{A} x_{G3C4} - K_{G3C5} x_{G3C5}\right) \tag{4.2.42}$$
$$v_{G3r6} = \kappa_{G3r6} \left(K_{A} K_{G3C5} x_{A} x_{G3C5} - K_{G3C6} x_{G3C6}\right) \tag{4.2.43}$$
$$v_{G3r7} = \kappa_{G3r7} \left(K_{A} K_{G3C6} x_{A} x_{G3C6} - K_{G3C7} x_{G3C7}\right) \tag{4.2.44}$$
$$v_{G3r8} = \kappa_{G3r8} \left(K_{A} K_{G3C7} x_{A} x_{G3C7} - K_{G3C8} x_{G3C8}\right) \tag{4.2.45}$$
$$v_{G3rt} = \kappa_{G3rt} \left(K_{G3C8} x_{G3C8} - K_{P3} K_{R} x_{P3} x_{R}\right) \tag{4.2.46}$$
$$v_{G3degM} = \kappa_{G3degM} \left(K_{G3M} x_{G3M} - K_{G3XM} x_{G3XM}\right) \tag{4.2.47}$$
$$v_{G3degP} = \kappa_{G3degP} \left(- K_{G3XP} x_{G3XP} + K_{P3} x_{P3}\right) \tag{4.2.48}$$

#### 4.3 List of State Derivatives $\dot{x}$ as Function of Flows $v$.

[15]:
```
## State equations
eqns = st.sprintdxl(s,sc,split=split)
disp.Latex(eqns)
```

[15]:

$$\dot{x}_{G1E} = -v_{G1Tc1} + v_{G1Tc2} - v_{G1Tc3} \tag{4.3.1}$$
$$\dot{x}_{G1EA} = v_{G1Tc1} - v_{G1Tc2} - v_{G1Tc4} \tag{4.3.2}$$
$$\dot{x}_{G1M} = v_{G1Tc2} - v_{G1rb} + v_{G1r1} - v_{G1degM} \tag{4.3.3}$$
$$\dot{x}_{P2} = -2v_{G1Tc3} - 2v_{G1Tc4} + v_{G2rt} - v_{G2degP} \tag{4.3.4}$$
$$\dot{x}_{G1EI} = v_{G1Tc3} \tag{4.3.5}$$
$$\dot{x}_{G1EAI} = v_{G1Tc4} \tag{4.3.6}$$
$$\dot{x}_{R} = -v_{G1rb} + v_{G1rt} - v_{G2rb} + v_{G2rt} - v_{G3rb} + v_{G3rt} \tag{4.3.7}$$
$$\dot{x}_{G1C0} = v_{G1rb} - v_{G1r1} \tag{4.3.8}$$
$$\dot{x}_{G1C1} = v_{G1r1} - v_{G1r2} \tag{4.3.9}$$
$$\dot{x}_{G1C2} = v_{G1r2} - v_{G1r3} \tag{4.3.10}$$

$$\dot{x}_{G1C3} = v_{G1r3} - v_{G1r4} \tag{4.3.11}$$
$$\dot{x}_{G1C4} = v_{G1r4} - v_{G1r5} \tag{4.3.12}$$
$$\dot{x}_{G1C5} = v_{G1r5} - v_{G1r6} \tag{4.3.13}$$
$$\dot{x}_{G1C6} = v_{G1r6} - v_{G1r7} \tag{4.3.14}$$
$$\dot{x}_{G1C7} = v_{G1r7} - v_{G1r8} \tag{4.3.15}$$
$$\dot{x}_{G1C8} = v_{G1r8} - v_{G1rt} \tag{4.3.16}$$
$$\dot{x}_{P1} = v_{G1rt} - v_{G1degP} - 2v_{G3Tc3} - 2v_{G3Tc4} \tag{4.3.17}$$
$$\dot{x}_{G2E} = -v_{G2Tc1} + v_{G2Tc2} - v_{G2Tc3} \tag{4.3.18}$$
$$\dot{x}_{G2EA} = v_{G2Tc1} - v_{G2Tc2} - v_{G2Tc4} \tag{4.3.19}$$
$$\dot{x}_{G2M} = v_{G2Tc2} - v_{G2rb} + v_{G2r1} - v_{G2degM} \tag{4.3.20}$$

$$\dot{x}_{P3} = -2v_{G2Tc3} - 2v_{G2Tc4} + v_{G3rt} - v_{G3degP} \tag{4.3.21}$$
$$\dot{x}_{G2EI} = v_{G2Tc3} \tag{4.3.22}$$
$$\dot{x}_{G2EAI} = v_{G2Tc4} \tag{4.3.23}$$
$$\dot{x}_{G2C0} = v_{G2rb} - v_{G2r1} \tag{4.3.24}$$
$$\dot{x}_{G2C1} = v_{G2r1} - v_{G2r2} \tag{4.3.25}$$
$$\dot{x}_{G2C2} = v_{G2r2} - v_{G2r3} \tag{4.3.26}$$
$$\dot{x}_{G2C3} = v_{G2r3} - v_{G2r4} \tag{4.3.27}$$
$$\dot{x}_{G2C4} = v_{G2r4} - v_{G2r5} \tag{4.3.28}$$
$$\dot{x}_{G2C5} = v_{G2r5} - v_{G2r6} \tag{4.3.29}$$
$$\dot{x}_{G2C6} = v_{G2r6} - v_{G2r7} \tag{4.3.30}$$

$$\dot{x}_{G2C7} = v_{G2r7} - v_{G2r8} \tag{4.3.31}$$
$$\dot{x}_{G2C8} = v_{G2r8} - v_{G2rt} \tag{4.3.32}$$
$$\dot{x}_{G3E} = -v_{G3Tc1} + v_{G3Tc2} - v_{G3Tc3} \tag{4.3.33}$$
$$\dot{x}_{G3EA} = v_{G3Tc1} - v_{G3Tc2} - v_{G3Tc4} \tag{4.3.34}$$
$$\dot{x}_{G3M} = v_{G3Tc2} - v_{G3rb} + v_{G3r1} - v_{G3degM} \tag{4.3.35}$$
$$\dot{x}_{G3EI} = v_{G3Tc3} \tag{4.3.36}$$
$$\dot{x}_{G3EAI} = v_{G3Tc4} \tag{4.3.37}$$
$$\dot{x}_{G3C0} = v_{G3rb} - v_{G3r1} \tag{4.3.38}$$
$$\dot{x}_{G3C1} = v_{G3r1} - v_{G3r2} \tag{4.3.39}$$
$$\dot{x}_{G3C2} = v_{G3r2} - v_{G3r3} \tag{4.3.40}$$

$$\dot{x}_{G3C3} = v_{G3r3} - v_{G3r4} \tag{4.3.41}$$
$$\dot{x}_{G3C4} = v_{G3r4} - v_{G3r5} \tag{4.3.42}$$
$$\dot{x}_{G3C5} = v_{G3r5} - v_{G3r6} \tag{4.3.43}$$
$$\dot{x}_{G3C6} = v_{G3r6} - v_{G3r7} \tag{4.3.44}$$
$$\dot{x}_{G3C7} = v_{G3r7} - v_{G3r8} \tag{4.3.45}$$
$$\dot{x}_{G3C8} = v_{G3r8} - v_{G3rt} \tag{4.3.46}$$

#### 4.4 List of Parameters.

[16]:
```
## Parameters
pars = st.sprintparl(parameter)
disp.Latex(pars)
```

[16]:

$$K_A = 485.2 \tag{4.4.1}$$
$$K_{G1E} = 1 \tag{4.4.2}$$
$$K_{G1EA} = 366.2 \tag{4.4.3}$$
$$K_{G1M} = 1 \tag{4.4.4}$$
$$K_{P2} = 4.54e-05 \tag{4.4.5}$$
$$K_{G1EI} = 9.358e-10 \tag{4.4.6}$$
$$K_{G1EAI} = 3.427e-07 \tag{4.4.7}$$
$$K_R = 1 \tag{4.4.8}$$
$$K_{G1C0} = 1 \tag{4.4.9}$$
$$K_{G1C1} = 3.49 \tag{4.4.10}$$

$$K_{G1C2} = 12.18 \quad (4.4.11)$$
$$K_{G1C3} = 42.52 \quad (4.4.12)$$
$$K_{G1C4} = 148.4 \quad (4.4.13)$$
$$K_{G1C5} = 518 \quad (4.4.14)$$
$$K_{G1C6} = 1808 \quad (4.4.15)$$
$$K_{G1C7} = 6311 \quad (4.4.16)$$
$$K_{G1C8} = 2.203e + 04 \quad (4.4.17)$$
$$K_{P1} = 4.54e - 05 \quad (4.4.18)$$
$$K_{G1XM} = 1e - 12 \quad (4.4.19)$$
$$K_{G1XP} = 1e - 16 \quad (4.4.20)$$

$$K_{G2E} = 1 \quad (4.4.21)$$
$$K_{G2EA} = 366.2 \quad (4.4.22)$$
$$K_{G2M} = 1 \quad (4.4.23)$$
$$K_{P3} = 4.54e - 05 \quad (4.4.24)$$
$$K_{G2EI} = 9.358e - 10 \quad (4.4.25)$$
$$K_{G2EAI} = 3.427e - 07 \quad (4.4.26)$$
$$K_{G2C0} = 1 \quad (4.4.27)$$
$$K_{G2C1} = 3.49 \quad (4.4.28)$$
$$K_{G2C2} = 12.18 \quad (4.4.29)$$
$$K_{G2C3} = 42.52 \quad (4.4.30)$$

$$K_{G2C4} = 148.4 \quad (4.4.31)$$
$$K_{G2C5} = 518 \quad (4.4.32)$$
$$K_{G2C6} = 1808 \quad (4.4.33)$$
$$K_{G2C7} = 6311 \quad (4.4.34)$$
$$K_{G2C8} = 2.203e + 04 \quad (4.4.35)$$
$$K_{G2XM} = 1e - 12 \quad (4.4.36)$$
$$K_{G2XP} = 1e - 16 \quad (4.4.37)$$
$$K_{G3E} = 1 \quad (4.4.38)$$
$$K_{G3EA} = 366.2 \quad (4.4.39)$$
$$K_{G3M} = 1 \quad (4.4.40)$$

$$K_{G3EI} = 9.358e-10 \quad (4.4.41)$$
$$K_{G3EAI} = 3.427e-07 \quad (4.4.42)$$
$$K_{G3C0} = 1 \quad (4.4.43)$$
$$K_{G3C1} = 3.49 \quad (4.4.44)$$
$$K_{G3C2} = 12.18 \quad (4.4.45)$$
$$K_{G3C3} = 42.52 \quad (4.4.46)$$
$$K_{G3C4} = 148.4 \quad (4.4.47)$$
$$K_{G3C5} = 518 \quad (4.4.48)$$
$$K_{G3C6} = 1808 \quad (4.4.49)$$
$$K_{G3C7} = 6311 \quad (4.4.50)$$

$$K_{G3C8} = 2.203e+04 \quad (4.4.51)$$
$$K_{G3XM} = 1e-12 \quad (4.4.52)$$
$$K_{G3XP} = 1e-16 \quad (4.4.53)$$
$$\kappa_{G1Tc1} = 30.85 \quad (4.4.54)$$
$$\kappa_{G1Tc2} = 0.0113 \quad (4.4.55)$$
$$\kappa_{G1Tc3} = 1e+06 \quad (4.4.56)$$
$$\kappa_{G1Tc4} = 1e+06 \quad (4.4.57)$$
$$\kappa_{G1rb} = 0.01 \quad (4.4.58)$$
$$\kappa_{G1r1} = 1.039e-05 \quad (4.4.59)$$
$$\kappa_{G1r2} = 2.976e-06 \quad (4.4.60)$$

$$\kappa_{G1r3} = 8.527e-07 \quad (4.4.61)$$
$$\kappa_{G1r4} = 2.443e-07 \quad (4.4.62)$$
$$\kappa_{G1r5} = 7e-08 \quad (4.4.63)$$
$$\kappa_{G1r6} = 2.005e-08 \quad (4.4.64)$$
$$\kappa_{G1r7} = 5.746e-09 \quad (4.4.65)$$
$$\kappa_{G1r8} = 1.646e-09 \quad (4.4.66)$$
$$\kappa_{G1rt} = 22.88 \quad (4.4.67)$$
$$\kappa_{G1degM} = 0.3466 \quad (4.4.68)$$
$$\kappa_{G1degP} = 3817 \quad (4.4.69)$$
$$\kappa_{G2Tc1} = 30.85 \quad (4.4.70)$$

$$\kappa_{G2Tc2} = 0.0113 \tag{4.4.71}$$
$$\kappa_{G2Tc3} = 1e+06 \tag{4.4.72}$$
$$\kappa_{G2Tc4} = 1e+06 \tag{4.4.73}$$
$$\kappa_{G2rb} = 0.01 \tag{4.4.74}$$
$$\kappa_{G2r1} = 1.039e-05 \tag{4.4.75}$$
$$\kappa_{G2r2} = 2.976e-06 \tag{4.4.76}$$
$$\kappa_{G2r3} = 8.527e-07 \tag{4.4.77}$$
$$\kappa_{G2r4} = 2.443e-07 \tag{4.4.78}$$
$$\kappa_{G2r5} = 7e-08 \tag{4.4.79}$$
$$\kappa_{G2r6} = 2.005e-08 \tag{4.4.80}$$

$$\kappa_{G2r7} = 5.746e-09 \tag{4.4.81}$$
$$\kappa_{G2r8} = 1.646e-09 \tag{4.4.82}$$
$$\kappa_{G2rt} = 22.88 \tag{4.4.83}$$
$$\kappa_{G2degM} = 0.3466 \tag{4.4.84}$$
$$\kappa_{G2degP} = 3817 \tag{4.4.85}$$
$$\kappa_{G3Tc1} = 30.85 \tag{4.4.86}$$
$$\kappa_{G3Tc2} = 0.0113 \tag{4.4.87}$$
$$\kappa_{G3Tc3} = 1e+06 \tag{4.4.88}$$
$$\kappa_{G3Tc4} = 1e+06 \tag{4.4.89}$$
$$\kappa_{G3rb} = 0.01 \tag{4.4.90}$$

$$\kappa_{G3r1} = 1.039e-05 \tag{4.4.91}$$
$$\kappa_{G3r2} = 2.976e-06 \tag{4.4.92}$$
$$\kappa_{G3r3} = 8.527e-07 \tag{4.4.93}$$
$$\kappa_{G3r4} = 2.443e-07 \tag{4.4.94}$$
$$\kappa_{G3r5} = 7e-08 \tag{4.4.95}$$
$$\kappa_{G3r6} = 2.005e-08 \tag{4.4.96}$$
$$\kappa_{G3r7} = 5.746e-09 \tag{4.4.97}$$
$$\kappa_{G3r8} = 1.646e-09 \tag{4.4.98}$$
$$\kappa_{G3rt} = 22.88 \tag{4.4.99}$$
$$\kappa_{G3degM} = 0.3466 \tag{4.4.100}$$

$$\kappa_{G3degP} = 3817 \tag{4.4.101}$$
